# Imaging minimal bacteria at the nanoscale: a reliable and versatile process to perform Single Molecule Localization Microscopy in mycoplasmas

**DOI:** 10.1101/2022.02.02.478797

**Authors:** Fabien Rideau, Audrey Villa, Pauline Belzanne, Emeline Verdier, Eric Hosy, Yonathan Arfi

## Abstract

Mycoplasmas are the smallest free-living organisms. These bacteria are important models for both fundamental and Synthetic Biology, owing to their highly reduced genomes. They are also relevant in the medical and veterinary fields, as they are pathogenic of both humans and most livestock species. Mycoplasma cells have minute sizes, often in the 300-800 nanometers range. As these dimensions are close to the diffraction limit of visible light, fluorescence imaging in mycoplasmas is often poorly informative. Recently developed “Super-Resolution Imaging” techniques can break this diffraction limit, improving the imaging resolution by an order of magnitude and offering a new nanoscale vision of the organization of these bacteria. These techniques have however not been applied to mycoplasmas before. Here, we describe an efficient and reliable protocol to perform Single-Molecule Localization Microscopy (SMLM) imaging in mycoplasmas. We provide a polyvalent transposon-based system to express the photo-convertible fluorescent protein mEos3.2, enabling Photo-Activated Localization Microscopy (PALM) in most *Mycoplasma* species. We also describe the application of direct STochastic Optical Reconstruction Microscopy (dSTORM). We showcase the potential of these techniques by studying the subcellular localization of two proteins of interest. Our work highlights the benefits of state-of-the-art microscopy techniques for mycoplasmology and provides an incentive to further the development SMLM strategies to study these organisms in the future.

**Importance:** Mycoplasmas are important models in biology, as well as highly problematic pathogens in the medical and veterinary fields. The very small size of these bacteria, well below the micron, limits the usefulness of traditional fluorescence imaging methods as their resolution limit is similar to the dimensions of the cells. Here, to bypass this issue, we established a set of state-of-the-art “Super-Resolution Microscopy” techniques in a wide range of *Mycoplasma* species. We describe two strategies: PALM, based on the expression of a specific photo-convertible fluorescent protein; and dSTORM, based on fluorophore-coupled antibody labeling. With these methods, we successfully performed single-molecule imaging of proteins of interest at the surface of the cells and in the cytoplasm, at lateral resolutions well below 50 nanometers. Our work paves the way toward a better understanding of mycoplasma’s biology through imaging of subcellular structures at the nanometer scale.

## Introduction

The colloquial term “mycoplasmas” refers to a set of bacteria belonging to the *Mollicutes* Class. These organisms derive from a common ancestor with Firmicutes through a degenerative evolution that has led to an extreme reduction in genome size (~0.6-1.35 Mbp). During this process, mycoplasmas have lost a large number of genes coding for important pathways, resulting in their characteristic lack of a cell wall and limited metabolic capacities (1–3). Owing to these deficiencies, mycoplasmas are obligate parasites that rely on their hosts for the production of a large array of essential metabolites. They have been isolated from a wide range of animals, including humans, mammals, reptiles, fish and arthropods.

Mycoplasmas are the simplest self-replicating organisms known to date and are thought to be good representatives of a “minimal” cell (4–6). They are therefore extremely interesting models in fundamental biology and have been used extensively to study the basic principles governing living systems and gene essentiality (7–11). These bacteria are also highly relevant in the field of Synthetic Biology, as their simplicity makes them prime models for the creation of engineered living systems. Mycoplasmas have been at the center of landmark studies such as the production of the first cell governed by a chemically synthesized genome, and later of the first minimal synthetic bacterial cell (12, 13). Mycoplasmas are also the first cells for which complete and accurate predictive mathematical models have been developed (14–16).

In parallel to these fundamental aspects, mycoplasmas are also highly problematic organisms in both the medical and veterinary fields as most of them are pathogenic for their hosts. In human, two species are particularly prevalent and concerning: *Mycoplasma genitalium,* which causes sexually transmissible urogenital infections (17); and *Mycoplasma pneumoniae* which causes “atypical pneumonia” predominantly in children and immunocompromised patients (18). While both pathogens typically cause mild diseases with low mortality, these infections are often chronic and the pathogens are not completely eliminated after antibiotics treatments (19, 20). Mycoplasmas also infect most livestock species and are major pathogens of cows (*Mycoplasma bovis*) (21), goats and sheeps (*Mycoplasma capricolum* subsp. *capripneumoniae*) (22), pigs (*Mycoplasma hyopneumoniae*) (23) and chickens (*Mycoplasma gallisepticum*) (24). Depending on the particular bacterial species or strain, mycoplasma infections can range from chronic, low mortality inflammatory diseases to peracute, highly lethal diseases. Infected animals often display heavily reduced production yields, endangering farmers’ revenues and threatening food security in poorly developed countries.

Studying mycoplasmas is a tedious process, as these organisms are slow growing and require complex and undefined culture media. In addition, the number of genetic tools available is limited, except for a small set of species belonging to the *Mycoides* cluster benefiting from techniques derived from the aforementioned Synthetic Biology projects (25).

The physical size of mycoplasmas is also a key limiting factor, as most species have cells with dimensions in the 300-800 nm range. These values are close to the resolution of diffraction-limited optical microscopy, which is in the 200–300 nm range with commonly used dyes and high numerical aperture oil immersion objectives. Thus, fluorescence microscopy in mycoplasmas is often poorly informative, as it is extremely difficult to determine the subcellular localization of the imaged component. This problem exists for most bacteria and archaea, and is exacerbated for mycoplasmas.

Higher resolution techniques based on immunogold labelling and electron microscopy have therefore been preferred to localize proteins at the cell surface or in the cytoplasm of mycoplasma cells (26–30). However, these methods suffer from complex sample preparation protocols, are difficult to set up for simultaneous visualization of multiple molecular species and are not compatible with live-cell imaging.

To this date, only a handful of studies have used immuno-fluorescence to study protein localization in mycoplasmas, and all of them have focused on ascertaining the polar distribution of proteins in the cells of *Mycoplasma mobile, M. pneumoniae* and *M. genitalium* which all exhibit highly polarized shapes (31–34). Similarly, a small number of studies have reported the polar localization or co-localization of proteins fused to the fluorescent proteins mCherry and EYFP, again in highly polarized cells (35–39). Other fluorescent proteins have successfully been expressed in several *Mycoplasma* species, including GFP (40), Venus (41), mNeonGreen, and mKO2 (42), but have only been used as expression reporters or transformation markers.

Interestingly, the last decade has seen the rapid development of multiple new fluorescence microscopy techniques aimed at bypassing the diffraction limit and bridging the gap between optical imaging resolution (~200 nm) and electron microscopy resolution (~1 nm). These methods, broadly termed “Single Molecule Localization Microscopy” (SMLM), rely on the successive imaging of individual fluorophores to mathematically determine their exact position (43–45). The spatiotemporal decorrelation of fluorescence emissions can be achieved in a stochastic manner, through the use of specific dyes or fluorescent proteins that can be made to randomly emit their fluorescence under wide-field illumination. Techniques such as Photo-Activated Localization Microscopy (PALM), Stochastic Optical Reconstruction Microscopy (STORM) or Points Accumulation for Imaging in Nanoscale Topography (PAINT) can typically yield images with lateral resolutions of ~10-50 nm. (46).

These methods have considerably expanded the possibilities of imaging in biology, allowing to resolve the subcellular organization of individual molecules or molecular assemblies, such as nuclear pores, chromatin complexes and cytoskeletal filaments at resolutions close to the molecular scale (47). The resolution improvements offered by SMLM appear especially attractive for microbiologists. These methods are have been progressively adopted by the scientific communities, first through their establishment in model bacteria such as *Escherichia coli* and *Bacillus subtilis*, and then by gradual transfer to more specialized fields (48–50). They have however not yet been applied to mycoplasmology.

In this report, we present the first protocols for the investigation of mycoplasmas using SMLM. We demonstrate the feasibility of PALM throughout the *Mollicutes* Class by successfully expressing a photo-convertible fluorescent protein in six highly relevant *Mycoplasma* species. Then, working in the model *Mycoplasma mycoides* subsp. *capri* (*Mmc*), we use PALM to image the subcellular localization of the cytoplasmic domain of an atypical F-type ATPase complex, yielding images with a lateral resolution of ~40 nanometers. In parallel, we apply dSTORM in the same model to study the localization of a surface-anchored protease involved in virulence, yielding images at a lateral resolution of ~25 nanometers.

## Material and Methods

### * Bacterial strains and culture conditions

*Escherichia coli* strain NEB-5α, used for plasmid cloning and propagation, was grown at 37°C in LB media supplemented with antibiotics (tetracycline 10 μg/mL; kanamycin 50 μg/mL; ampicillin 100 μg/mL) when selection was needed.

*Mycoplasma mycoides* subsp. *capri* strain GM12 (*Mmc*), *Mycoplasma capricolum* subsp. *capricolum* strain 27343 (*Mcap), Mycoplasma bovis* strain PG45 (*Mbov), Mycoplasma agalactiae* strain PG2 (*Maga*), *Mycoplasma genitalium* strain G37 (*Mgen*), *Mycoplasma gallisepticum* strain S6 (*Mgal*) were grown at 37°C in appropriate media: SP5 (51), Hayflick modified (52) or SP4 modified (52, 53), supplemented with antibiotics (gentamycin 100-400 μg/mL; tetracycline 5 μg/mL) when selection was needed.

### * Cloning the mEOS3.2 expression plasmids

The mycoplasma codon-optimized version of the coding sequence for mEos3.2 was designed using the online tool JCat (http://www.jcat.de/; parameters: input= protein sequence, reference organism= *Mycoplasma mycoides* (subsp. mycoides SC, strain PG1), options= “Avoid prokaryotic ribosome binding sites”). Further adaptation of the codon usage was performed manually based on the recommendation of the Twist Bioscience synthesis tool, to enable the chemical synthesis of the corresponding DNA fragment (Twist Bioscience). After PCR amplification, the coding sequence was cloned by InFusion (Clontech) in the plasmid pMT85 under the control of the promoter PTufA (42). Based on this pMT85-PTufA-mEos3.2 plasmid, three other variants were subsequently produced by Gibson Assembly (NEB) or by site-directed mutagenesis (NEB) to replace the original promoter by either the promoter PSpi (51), the promoter PSynMyco (54), or the promoter P438 (55)). Cloned plasmids were transformed in chemically competent *E. coli* NEB-5α for maintenance and propagation, then isolated by miniprep, verified by enzymatic restriction and finally checked by Sanger sequencing (Genewiz).

### * Mycoplasmas transformation

Plasmids pMT85-PSynMyco-mEos3.2, pMT85-PSpi-mEos3.2 or pMT85-P438-mEos3.2 were transformed by PolyEthylene Glycol contact in *Mmc*, *Mcap* (56), *Maga* and *Mbov* (57) and *Mgal* (unpublished); or by electroporation in *Mgen* (58). Transformants were subsequently plated on appropriate media plates containing antibiotics for selection (see above). Transformants clones were then passaged 3 times in liquid media supplemented with gentamycin (100-400 μg/mL). The presence of the mEOS3.2 coding sequence was checked by PCR.

### * Production of Mycoplasma mycoides subsp. mycoides mutants

Accurate edition of the genome of *Mmc* is currently only possible through a complex process involving the transfer of the bacterial chromosome in a yeast cell, its modification in the yeast, and its subsequent transplantation back in a *Mcap* bacterial recipient cell (59). This process was used to generate two mutant strains: *Mmc* 0582-HA in which an HA-tag coding sequence is fused to the C-terminus of the coding sequence of MMCAP2_0582; and *Mmc* mEos3.2-0575 in which the codon-optimized mEos3.2 coding sequence is fused to the N-terminus of the coding sequence of MMCAP2_0575. To do so, we first generated two plasmids encoding guide RNAs targeting either MMCAP2_0582 or MMCAP2_0575 by modification of the base plasmid p426-SNR52p-gRNA.Y-SUP4t (25) using the Q5 Site-Directed Mutagenesis kit (NEB). In addition, we generated two recombination cassettes that contain the modified version of the target loci flanked by 1000 bp recombination arms. Both cassettes were built by first cloning the wild-type locus and 1 kb flanking region in the pGEM-T plasmid (Promega) and subsequently modifying the plasmid to add the HA-tag coding sequence using the Q5 Site-Directed Mutagenesis kit (NEB) or the mEos3.2 coding sequence using InFusion (Clontech). Plasmids carrying the cassettes were checked by Sanger sequencing (Genewiz), and the cassettes amplified by PCR. Both the guide RNA-encoding plasmid and the recombination cassette were then co-transformed in the yeast carrying the *Mmc* chromosome. Yeast transformants were checked for the presence of the integral bacterial chromosome by multiplex PCR and for the presence of the desired mutation by PCR and amplicon sequencing. The modified genome was then extracted and transplanted in the recipient cell. The resulting transplants were also checked for genome integrity by multiplex PCR and for the presence of the desired mutation PCR and amplicon sequencing.

### * Sample preparation for SMLM experiments

Mycoplasmas cells, grown to late log phase at 37°C in appropriate media supplemented with gentamycin (100-400 μg/mL), were harvested by centrifugation at 6800 rcf, 10°C for 10 minutes. After removal of the spent media, the cells were washed twice in one volume of buffer (HEPES 67.7 mM - NaCl 140 mM - MgCl2 7 mM - pH 7.35), and subsequently resuspended in 1/3^rd^ volume of the same buffer. When necessary, cell aggregates were broken down by performing 20 passages through a 26 Gauge needle. A poly-L-lysine coated 1.5H 18 mm precision coverslip (Marienfeld) was placed at the bottom of a 12-well plate and equilibrated in 1 mL of wash buffer. Three microliters of the cell suspension were added to the well, and the plate was subsequently centrifuged in a swing-out rotor at 2500 rcf, 10°C for 10 minutes to force the cells to sediment on the coverslip. The well was emptied by suction, and the coverslip was then moved to a clean well. The coverslip was then washed once with 3 mL wash buffer and then incubated in 1 mL of a pre-warmed 4% PFA in washing buffer, at 37°C for 30 min. Five washing steps with 3 mL of wash buffer were performed to eliminate all PFA traces. Correct deposition and fixation of the cells on the coverslip was checked by dark-field microscopy (Nikon Eclipse Ni microscope equipped with a dark field condenser and a Teledyne Photometrics Iris 9 camera) using washing buffer as mounting medium. Validated coverslips were stored at 4°C in PBS until ready to use for PALM imaging (no more than 10 days).

For dSTORM imaging, coverslips were prepared as described above. After the final wash, each coverslip was incubated 1 hour at room temperature in 1 mL of blocking buffer (PBS/BSA 1%). Immuno-labeling was performed by first using a mouse anti HA-tag primary antibody (Thermo) diluted at 1/500^th^ in blocking buffer. After incubation for 1 hour at room temperature, the coverslip was washed three times with 1 mL of PBS. Then, the coverslip was incubated for 1 hour at room temperature with the secondary antibody (goat anti-mouse IgG conjugated to the Alexa647; Jackson ImmunoResearch) diluted at 1/500^th^ in 1 mL of blocking buffer. After three wash steps in PBS, an additional fixation was performed by incubating the labeled coverslips with 2% PFA in PBS for 5 minutes at 37°C. Finally, the coverslips were washed five times in wash buffer and stored at 4°C in PBS until imaging (no more than 10 days).

### * SMLM imaging set up and data collection

Imaging was performed on a LEICA DMi8 inverted microscope, mounted on an anti-vibrational table (TMC, USA), using a Leica HC PL APO 100 × 1.47 NA oil immersion TIRF objective and fiber-coupled laser launch (405 nm, 488 nm, 532 nm, 561 nm and 642 nm) (Roper Scientific, Evry, France). Fluorescence signal was collected with a sensitive Evolve EMCCD camera (Teledyne Photometrics). The coverslips bearing the fixed bacterial cells were mounted on a Ludin chamber (Life Imaging Services) and 600 μL of imaging buffer was added. For PALM imaging, the buffer was PBS. For dSTORM imaging, the buffer contained both oxygen scavenger (glucose oxidase) and reducing agent (2-Mercaptoethylamine), and another 18 mm coverslip was added on top of the chamber to minimize oxygen exchanges during the acquisition.

Image acquisition and control of the microscope were driven through Metamorph (Molecular devices) in streaming mode using a 512 x 512 pixels region of interest with a pixel size of 160 nm. Image stacks typically contained 6,000–20,000 frames, acquired at a frequency of 33 Hz for PALM and 50 Hz for dSTORM. The power of the 405 nm laser was adjusted to control the density of single molecules per frame, keeping the 642 nm laser intensity constant. For dual-color imaging, dSTORM was performed first followed by PALM. To limit manipulation and the potential resulting drift, coverslips were kept in dSTORM imaging medium during the PALM acquisition.

### * SMLM data processing and analysis

The PALMTracer plugin for Metamorph (60–62) was used to process image stacks with a specific intensity threshold for each dataset, to enable the generation of the localization tables. From these tables, super-resolved images were generated with a pixel dimension of 40 nm. Pointing accuracies of the PALM imaging experiments were determined by tracking individual fluorophore’s motion using PALMTracer. For each track with a speed below 0.01 μm^2^.s^−1^, the half of the root of the MSD0 value was calculated (63). PALM experiments are done with a pointing accuracy of 41 nm in average). Resolution of dSTORM imaging experiments was determined by Gaussian fitting of the signal of individual fluorophores attached to the coverslip. Both resolution (66 nm) and pointing accuracy (25 nm) are extracted.

Clustering of the localizations was performed by Voronoï segmentation using SR-Tesseler (64). Object detection and clustering characterization were done using a density factor of 1, and a minimum cluster area of 0.05 pixel^2^. The cell-counter plugin of Fiji was used to count the number of cells present in our acquisition field. Dot plots were graphed using PlotsOfData (65).

## Results

### 1. Establishing a common and efficient sample preparation process for SMLM imaging in mycoplasmas

The production of high-quality samples is generally regarded as a critical step for the acquisition of high-quality SMLM data, and multiple reviews provide important guidelines to follow (66, 67). Here, the first step of the process was to ensure the reliable immobilization of the mycoplasma cells to high-precision glass coverslips. We initially attempted to grow the mycoplasmas directly onto poly-L-lysine coated coverslips by immersing the coverslips in inoculated media. However, this approach failed as cells remained planktonic and did not adhere to the glass. We thus developed an efficient and reliable centrifugation-based process to force the cells to sediment and attach to the coverslip (Figure S1). Briefly, mycoplasma cells are harvested and washed to form a homogenous medium-density suspension (approximately 10^5^-10^7^ cfu.mL^−1^) in which the coverslip is immersed, in a 12 well plate. Centrifugation at low speed in a swing-out rotor forces the cells to sediment and adhere to the glass. After washing to remove unbound cells, fixation is performed using a solution of 4% paraformaldehyde. The coverslip quality is checked by observation with a dark-field microscope. A good sample is characterized by bacterial cells deposited in a monolayer, regularly spaced and separated from each other by a few microns. These samples can either be imaged directly using PALM or further processed by immuno-labeling in order to later perform dSTORM.

### 2. Establishing a polyvalent mEos3.2 expression system for PALM in multiple mycoplasmas

PALM imaging is based on the expression of specific fluorescent proteins that are either photoactivatable (irreversible off-to-on) or photo-convertible (wavelength A to wavelength B) (68). This process is driven by light, and can be tuned to occur at a slow rate, thus enabling imaging of individual fluorophores. Here, we elected to use the fluorescent protein mEos3.2 (69), which can be converted by illumination with a near-UV wavelength (405 nm) from a green state (Ex= 507 nm, Em= 516 nm) to a red state (Ex= 572 nm, Em= 580 nm).

To assess the functionality of mEos3.2 in a wide range of *Mycoplasma* species, we first designed a mEos3.2 codon-optimized coding sequence using the codon usage table of *Mycoplasma mycoides* subsp. *mycoides* strain PG1 as reference. This coding sequence was subsequently cloned in the plasmid pMT85 (42, 58, 70–74). The plasmid backbone carries a transposon derived from Tn4001 (75), the insertion of which can be selected through the gentamicin resistance gene *aacA-aphD* placed under the control of its natural promoter. In order to drive the expression of the mEos3.2, we used the recently developed synthetic regulatory region PSynMyco (54). These three elements (transposon, selection marker and SynMyco regulatory region) have all been shown to be functional in a wide range of *Mollicutes* species, and should yield a “mycoplasma-universal” mEos3.2 expression vector.

The resulting plasmid pMT85-PSynMyco-mEos3.2 (Figure 1A) was transformed in six *Mycoplasma* species, relevant to either the veterinary or medical fields, and covering the three main *Mollicutes* phylogenetic sub-groups (Figure 1B): *i*) *Mycoplasma mycoides* subsp. *capri* strain GM12 (*Mmc*) and *Mycoplasma capricolum* subsp. *capricolum* strain 27343 (*Mcap),* Spiroplasma group; *ii*) *Mycoplasma bovis* strain PG45 (*Mbov*) and *Mycoplasma agalactiae* strain PG2 (*Maga*), Hominis group; and *iii*) *Mycoplasma genitalium* strain G37 (*Mgen*) and *Mycoplasma gallisepticum* strain S (*Mgal*), Pneumoniae group. Transformants carrying the transposon were obtained for all six species, and sample coverslips were prepared for each.

**Figure 1:**
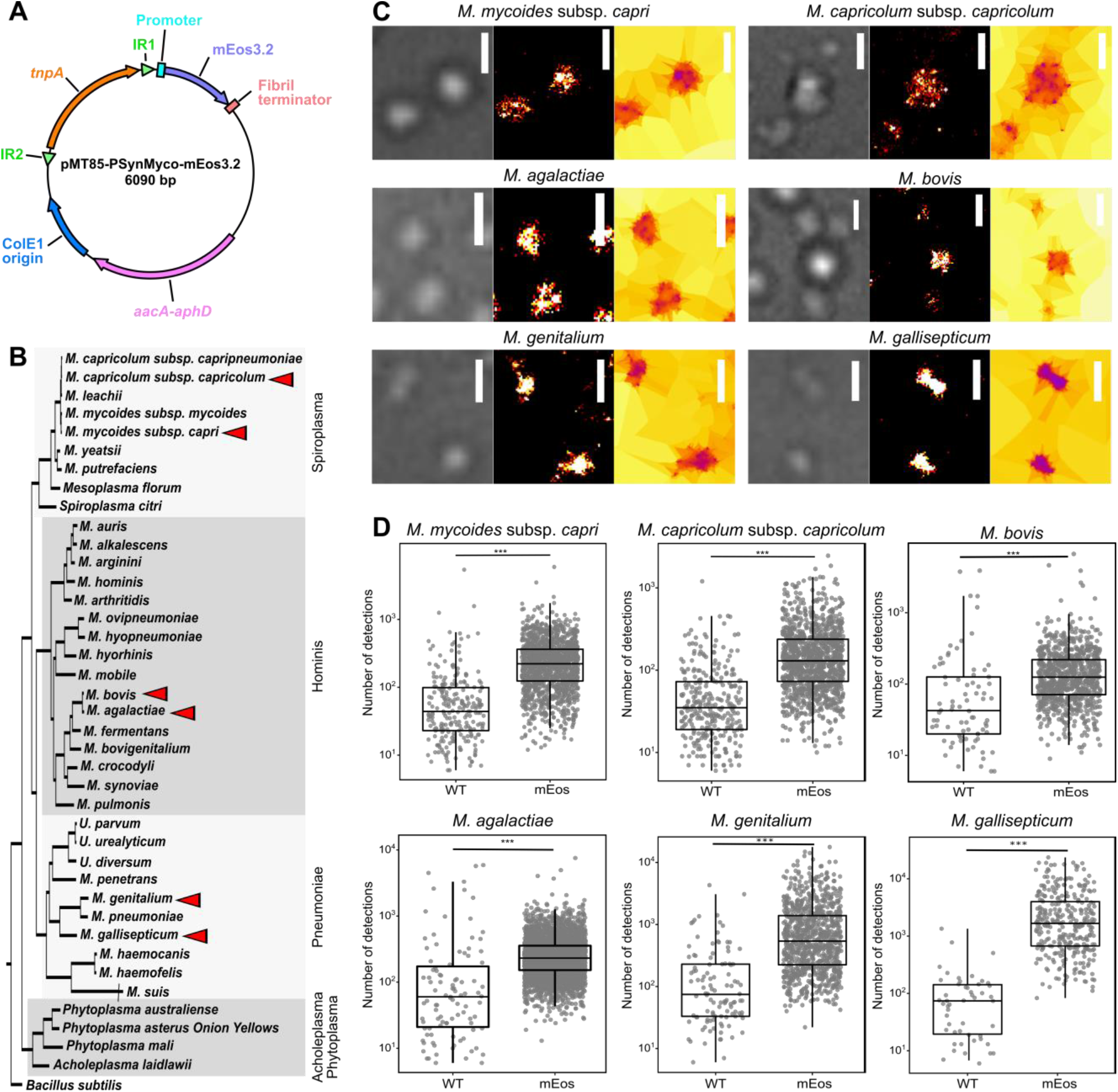
Assessing the functionality of the photo-convertible fluorescent protein mEos3.2 in multiple *Mycoplasma* species. **A.** Map of the plasmid pMT85-PSynMyco-mEos3.2. The main genetic components of the plasmid are indicated (IR: Inverted Repeat, *tnpA:* transposase, *aacA-aphD:* gentamycin resistance). **B.** Distribution of the *Mycoplasma* species used in this study. A phylogenetic tree of representative *Mollicutes* species was inferred using the maximum-likelihood method from the concatenated multiple alignments of 79 proteins encoded by genes present at one copy in each genome (adapted from Grosjean *et al*. 2014). The main phylogenetic groups are indicated by grey boxes. The six species transformed with pMT85-PSynMyco-mEos3.2 are identified by a red arrow. **C.** Sample images of mycoplasma cells expressing the mEos3.2 in their cytoplasm. For each of the six species, cells transformed with the plasmid pMT85-PSynMyco-mEos3.2 were imaged using PALM. A representative subset of the field of view is given (from left to right: contrast-phase image, super-resolved reconstruction at 40 nm pixels, Tesseler segmentation of the localizations). Scale bar: 1 μm. **D.** Quantification of the PALM signal intensity. For each of the six species, both wild-type (WT) and pMT85-PSynMyco-mEos3.2 transformed cells (mEos) were imaged by PALM and the data collected from a single representative field of view (512×512 pixels; pixel size: 0.16 μm) were analyzed. The dot-plot presents the number of detections measured in each object segmented by Tesseler (equivalent to a cell), on top of which a boxplot showing the median, interquartile range, minimum and maximum is overlaid. Statistics tests (Mann Whitney) were performed to compare the two conditions WT and mEos (***: p-value < 1.10^−10^).

We then assessed the ability of the mEos3.2 to be converted to its red state by performing PALM imaging. Cells were observed in the red wavelength, with low power illumination at 405 nm to sparsely and stochastically photo-convert the mEos3.2. Based on the acquired image stacks, the localization of each individual fluorescence emitter was determined through the PALMtracer plugin under Metamorph, then, super-resolved images are reconstructed and analyzed by automatic Voronoï-based segmentation of these localizations (Figure 1C, Figure S2). The principle of tessellation analysis is to extract objects with a similar density of localization. Here, with the first level of segmentation we isolated the property of individual mycoplasma cells, with the second level yielding data on clustering of the protein of interest.

For all the six *Mycoplasma* species studied, wild type and mutant, a comparable number of cells per field of view is observed in transmission light Figure S3A). For each first level segmented object, the number of underlying detections was compared between the wild-type and mEos3.2 expressing mycoplasmas. Similar results were obtained for each of the six *Mycoplasma* species studied (Figure 1D). For the wild-type cells, only a small number of individual objects were identified (*Mmc:* 240, *Mcap:* 291, *Mbov:* 79, *Maga:* 99, *Mgen:* 110, *Mgal:* 50), with each being supported by a small number of detections (median detections/object: *Mmc:* 42, *Mcap:* 31, *Mbov:* 43, *Maga:* 63, *Mgen:* 61, *Mgal:* 79). Amongst them, a large fraction of the objects were found outside of the cells, suggesting that they are imaging artifacts rather than signals emitted by the mEos3.2 (Figure S3B). Comparatively, the mEos3.2 expressing cells yielded one to two orders of magnitude more objects (*Mmc:* 1487, *Mcap:* 1183, *Mbov:* 814, *Maga:* 3761, *Mgen:* 919, *Mgal:* 326), each supported by one order of magnitude more detections (median detection/object *Mmc*: 245, *Mcap*: 132, *Mbov*: 152, *Maga*: 237, *Mgen*: 493, *Mgal*: 1826; Figure 1D). In the mEos samples, most objects are found inside the boundaries of the cells (Figure S3B). Taken together, the data collected indicate that the mEos3.2 is functional in a wide range of *Mycoplasma* species, thus enabling PALM imaging of proteins of interest in these organisms.

### 3. PALM imaging of an atypical F-ATPase in Mmc

To showcase the interest of performing PALM to study biological processes in mycoplasmas, we worked in our model organism *Mmc*. We were particularly interested in localizing an atypical F-type ATPase called “F1-like-X0” (76), putatively involved in a mechanism of immune evasion (59). Leveraging the genome engineering tools available for this species, we generated a mutant strain *Mmc* mEos3.2-0575 in which the fluorescent protein is fused to the N-terminus of the β-subunit of the F1-like domain. Diffraction-limited images taken by epifluorescence in the green wavelength (before photo-conversion) did not yield any meaningful information, as they only show dimly lit cells with no apparent distribution of the signal (Figure 2A). In stark contrast, the super-resolved images reconstructed from the PALM datasets reveal that most of the signal detected for each cell is localized in a small number of clusters that are found predominantly at the periphery of the cytoplasm (Figure 2A). This localization is in accordance with pre-existing information gathered on the association of the F-domain of this ATPase with the internal face of the plasma membrane (76). The first level of tessellation successfully delineated the cells, and automatic segmentation of the localizations indicated that each cell typically presents 2 to 4 clusters of detections (mean= 3.3, median= 3) (Figure 2B). Meanwhile, cells expressing mEos3.2 as a monomer in the cytoplasm presented mainly a single cluster (Figure S4). The median area of the clusters formed by mEos-tagged ATPase is 5609 nm^2^ (Figure 2C) which corresponds to a circle with a diameter of 84 nm. This cluster area corresponds on average to ~2% of the median cell area, estimated to 324518 nm^2^ (equivalent to a circle of 640 nm of diameter).

**Figure 2:**
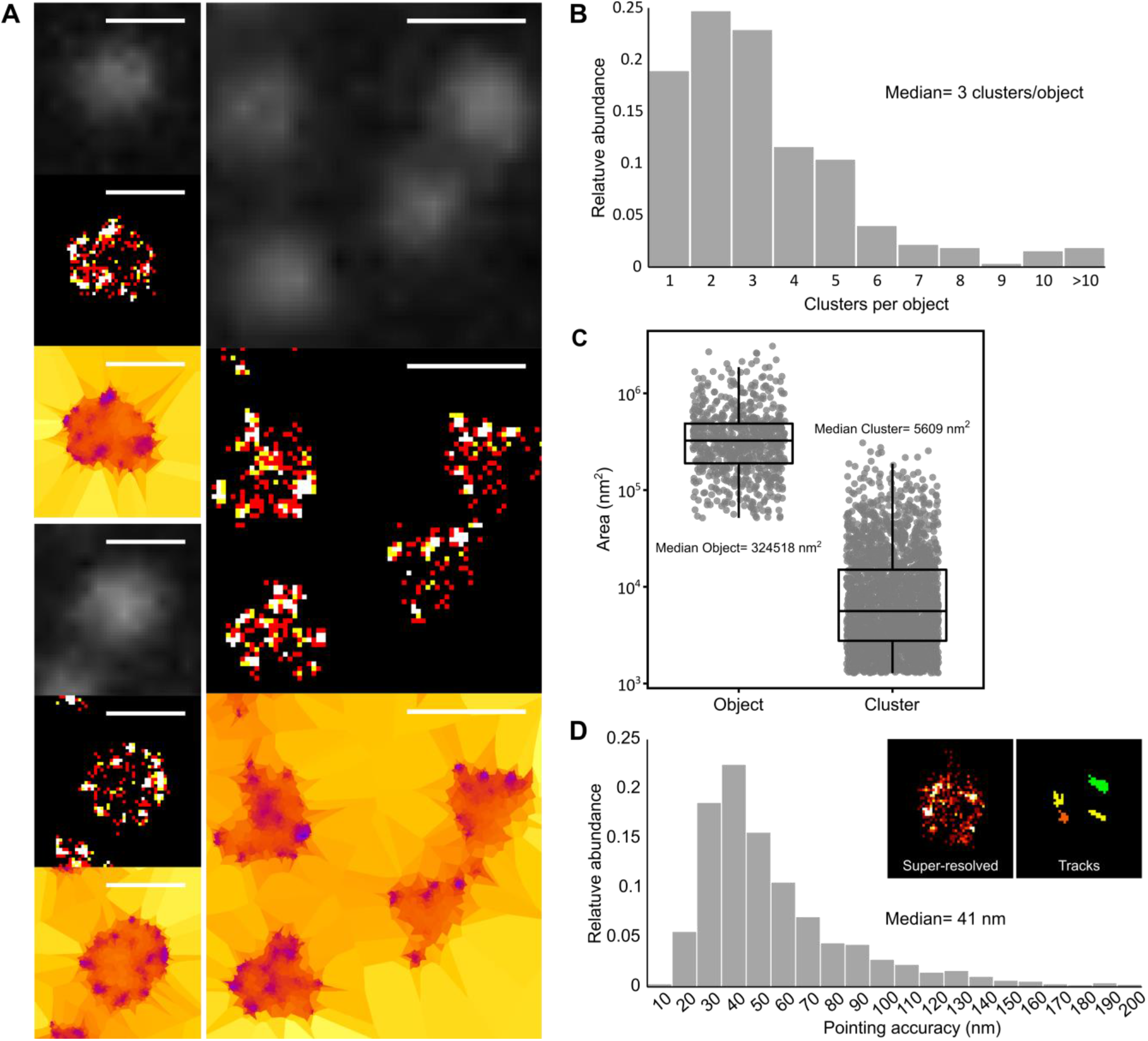
PALM imaging of an F-type ATPase in *Mycoplasma mycoides* subsp. *capri.* *Mmc* mEos3.2-0575 cells, expressing a mEos-fused variant of the β-subunit of the ATPase F1-like domain, were imaged by PALM. In this *Mmc* mutant, the fluorescent fusion protein is expressed from the native genomic locus and replaces the wild-type variant. The data presented here correspond to a single representative field of view (512×512 pixels; pixel size: 160 nm). **A.** Sample images of *Mmc* mEos3.2-0575 cells. For each field of view, the images correspond to: top: epifluorescence (diffraction-limited); middle: super-resolved reconstruction (40 nm pixel); bottom: Tesseler segmentation. Scale bar: 1 μm. **B.** Tesseler clustering of the fluorescence signal. The number of clusters per Tesseler-segmented objects was computed. The bar graphs display the distribution of the number of clusters per object. **C.** Objects and clusters sizes. The dot plot presents the area (in nm^2^) of each object and cluster segmented by Tesseler, to which a boxplot showing the median, interquartile range, minimum and maximum is overlaid. The median value of each data set is indicated. **D.** Evaluation of the PALM imaging pointing accuracy. Inset: example of the tracks computed using PALM-Tracer and from which the MSD0 and pointing accuracies values are derived. The bar graphs display the distribution of the pointing accuracies derived from each track. The median value of the data set is indicated.

In order to estimate the accuracy of our detections, we used PALM-tracer to track individual emitters and calculate the Mean squared displacement (MSD) of the trajectories (Figure 2D, inset). The fit of the first points of the MSD gives access to both the median speed of the molecule and its pointing accuracy (77). As our cells are fixed, we directly obtained the pointing accuracy of the fluorophore (63). The majority of pointing accuracies were between 30 and 60 nm, with a median of 41 nm (Figure 2D). The imaging process was highly reproducible, with similar results obtained for data collected from two regions of the same coverslip or from two independent coverslips (Figure S5: Median clusters per object: 3; Median object area: 329971-464727 nm^2^ – equivalent to a circle with a diameter of 648-770 nm; Median cluster area: 6188-8795 nm^2^ – equivalent to a circle with a diameter of 88-106 nm; Median pointing accuracy: 39-51 nm).

### 4. dSTORM imaging of a surface protease in Mmc

In parallel to PALM, we also performed dSTORM imaging in *Mmc.* In this method, a fluorophore-conjugated probe, such as an antibody, is used to label the protein of interest to image. The fluorophores used in dSTORM have the ability to spontaneously transition in and out of a dark state, under strong excitation illumination and in a reducing and oxygen-depleted buffer. To test this method in mycoplasma, we first generated a mutant strain of *Mmc* in which the HA epitope tag was fused to the C-terminus of the protease MIP0582 (59, 78). This immunoglobulin-specific protease is anchored to the cell surface and belongs to the same immune evasion system as the F-ATPase studied above. Coverslips samples of *Mmc* 0582-HA were immuno-labeled with a mouse anti-HA primary antibody and a goat anti-mouse Alexa647-conjugated secondary antibody. Acquisition of diffraction-limited data was performed by illuminating the sample with the excitation laser at a low power which is not sufficient to cause blinking of the fluorophores. Again, no meaningful information could be extracted as the cells appeared dimly fluorescent, with a slightly more intense signal at the periphery in some cases (Figure 3A). Conversely, super-resolved images reconstructed from the dSTORM datasets reveal that the fluorescence signal is localized in clusters that are predominantly at the periphery of the cell (Figure 3A). This is in accordance with the known localization of the protease at the cell surface.

**Figure 3:**
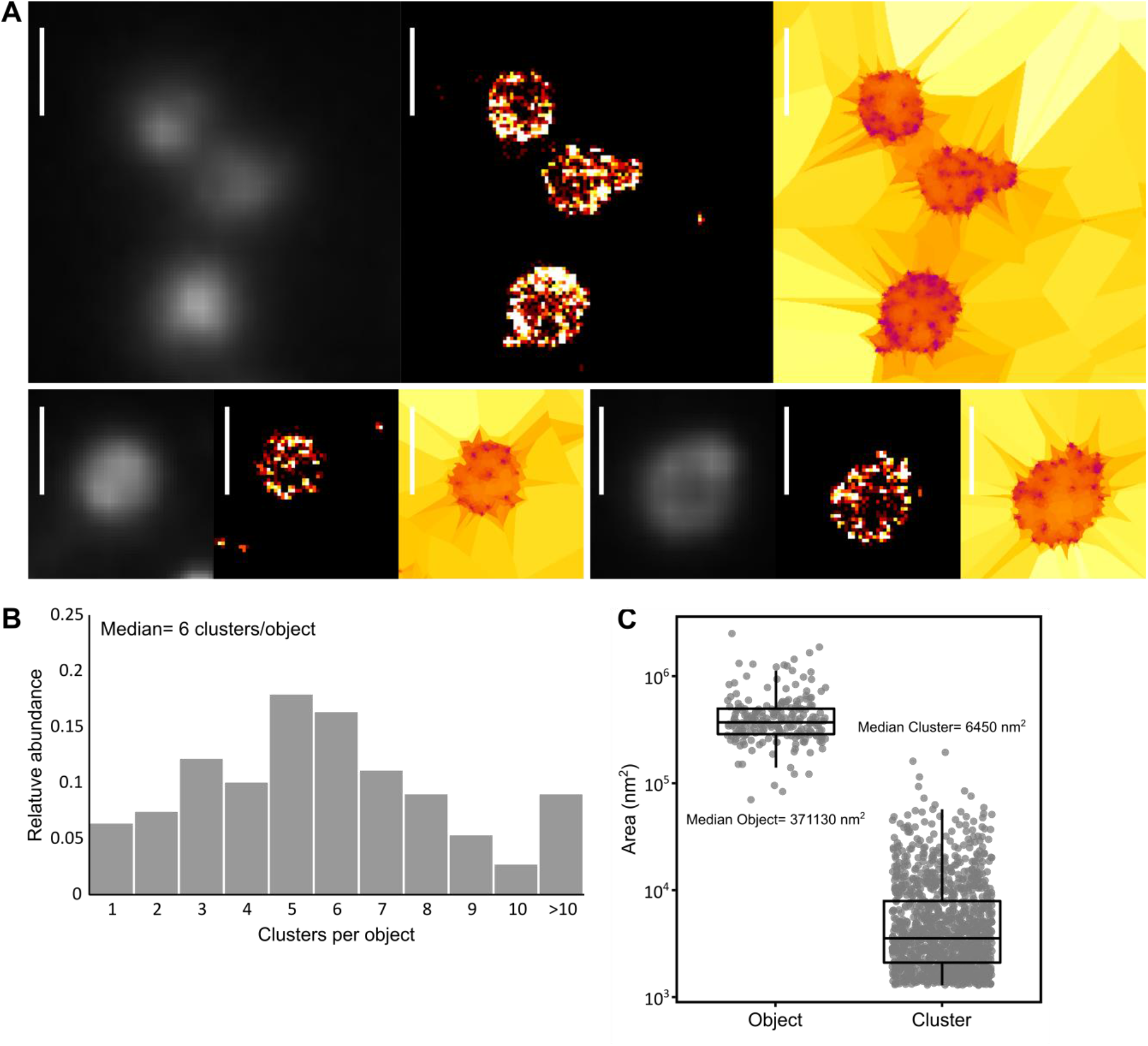
dSTORM imaging of an antibody-specific protease in *Mycoplasma mycoides* subsp. *capri*. *Mmc* 0582-HA cells expressing an HA-tag fused variant of the serine protease MIP_82_ were immune-labeled and imaged by dSTORM. The tagged fusion protein is expressed from the native genomic locus and replaces the wild-type variant. The data presented here correspond to a single representative field of view (512×512 pixels; pixel size: 160 nm). **A.** Sample images of *Mmc* 0582-HA cells. For each field of view, the images correspond to: left: epifluorescence (diffraction-limited); middle: super-resolved reconstruction (40 nm pixel); right: Tesseler segmentation. Scale bar: 1 μm. **B.** Tesseler clustering of the fluorescence signal. For each field of view, the number of clusters per Tesseler-segmented objects was computed. The bar graphs display the distribution of the number of clusters per object. **C.** Objects and clusters sizes. The dot plot presents the area (in nm^2^) of each object and cluster segmented by Tesseler, to which a boxplot showing the median, interquartile range, minimum and maximum is overlaid. The median value of each data set is indicated.

Automatic segmentation of these localizations indicates that cells typically present 5-10 clusters (median= 6), and a significant proportion of cells exhibit more than 10 clusters (10%) (Figure 3B). The median cluster area is 3254 nm^2^ (Figure 3C) which corresponds to a circle with a diameter of 64 nm. Interestingly, these clusters are smaller than those measured by PALM for the ATPase, with a median area inferior by ~40% and a corresponding circle diameter inferior by ~25%, probably due to the higher pointing accuracy obtained with dSTORM technique (25 nm) compared to PALM (41 nm). However, the first level of tessellation objects, which can be assimilated to the cells, are very similar to those observed by PALM, with a median area of 371130 nm^2^ (equivalent to a circle with a diameter of 687 nm). Again, the imaging process showed high reproducibility, with similar results from data collected from two regions of the same coverslip or from two independent coverslips (Figure S6: Median clusters per object: 5-7; Median object area: 353506-422145 nm^2^ – equivalent to a circle with a diameter of 670-733 nm; Median cluster area: 3582-3944 nm^2^ – equivalent to a circle with a diameter of 67-70 nm).

### 5. Dual-color and 3D SMLM in Mmc

One of the key advantages of fluorescence imaging is the ability to perform multi-color experiments to locate multiple proteins of interest by combining fluorophores with different excitation/emission spectra. Here, we leveraged the spectral compatibility between the mEos3.2 red state and the Alexa647 emissions/excitations wavelengths to perform dSTORM/PALM dual color imaging in *Mmc.* To do so, the strain *Mmc* 0582-HA was transformed with the plasmid pMT85-PSynMyco-mEos3.2. The transformants were then imaged sequentially, first by dSTORM, then by PALM. Datasets were processed as above. Super-resolved images were successfully reconstructed from both datasets, yielding results similar to those observed during single-color imaging: the mEos filling the cytoplasm in PALM, and 10-20 small clusters of MIP0582-HA at the cell periphery in dSTORM (Figure 4A). This experiments clearly demonstrate the ability of super-resolution technique to discriminate two types of organization in the minute mycoplasma cells, while classical microscopy failed.

**Figure 4:**
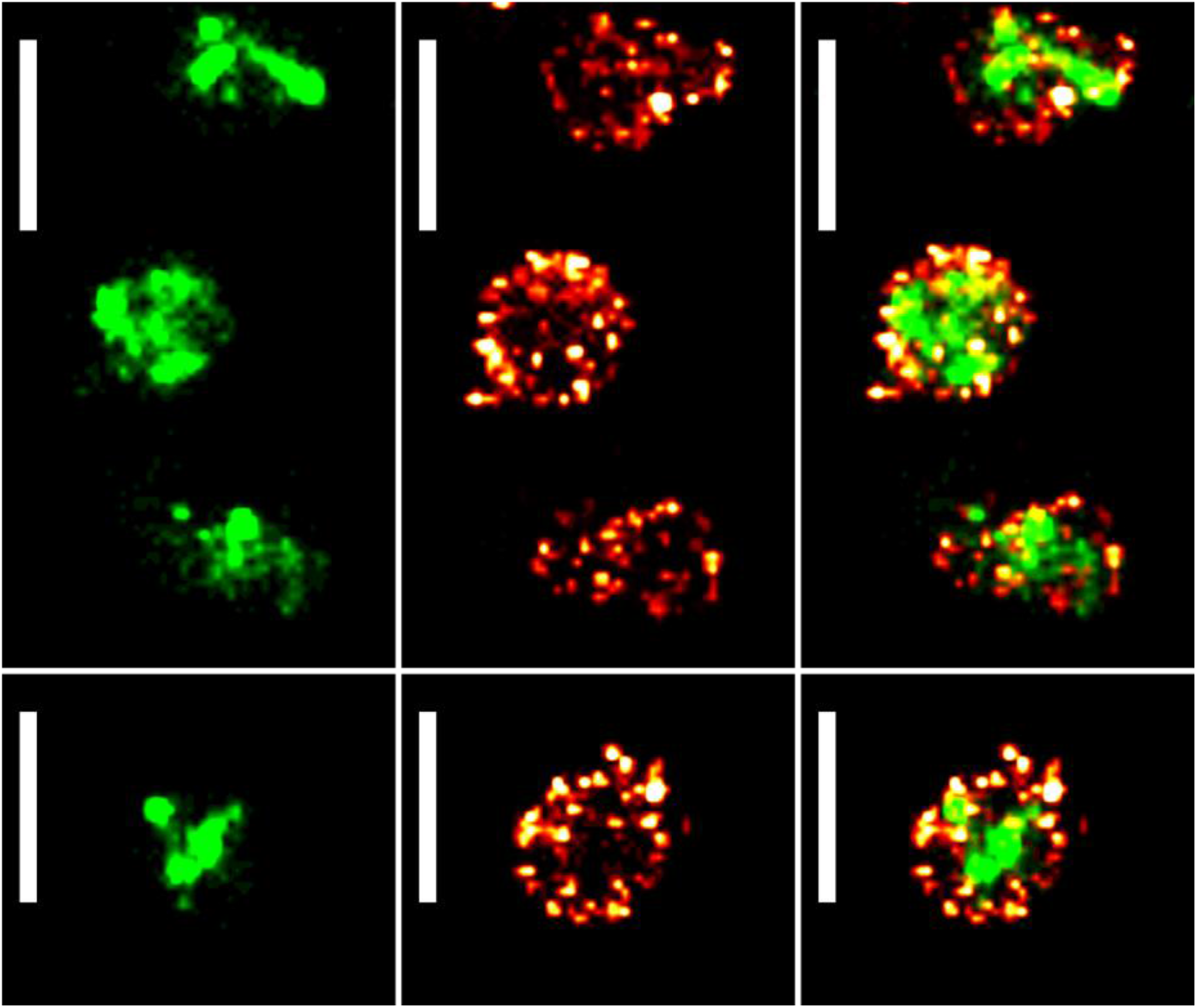
PALM/dSTORM two-colors imaging of *Mycoplasma mycoides* subsp. *capri.* Sample images of *Mmc* 0582-HA pMT85-PSynMyco-mEos3.2 cells, expressing both an HA-tag fused variant of the serine protease MIP0582 and the fluorescent protein mEos3.2. The tagged fusion protein is expressed from the native genomic locus and replaces the wild-type variant. The mEos3.2 is expressed from the transposon inserted at a random site in the bacterial chromosome. For each field of view, the images correspond to: left: reconstructed PALM image (40 nm pixel); middle: reconstructed dSTORM (40 nm pixel); right: overlay of the reconstructed PALM and dSTORM images. Scale bar: 1 μm. All the images were sampled from the same coverslip and field of view.

### 6. Using mEos3.2 as a reporter and PALM imaging to measure physical parameters

In addition to the acquisition of localization data for proteins of interest, we also demonstrated that PALM imaging process could be used to gather information on protein expression levels. Indeed, similarly to what can be done with classical fluorescent proteins, the amount of mEos3.2 in a given cell is proportional to the number of detections. We compared the relative strength of various promoters by building modified versions of the plasmid pMT85-PSynMyco-mEos3.2 in which the PSynMyco promoter was replaced by the P438 or PSpi promoter, respectively. The three plasmids were transformed in *Mmc*, yielding transformants that were subsequently imaged by PALM. Analysis of the data revealed that each promoter drives the expression of mEos3.2 at different levels (Figure 5A), with P438 producing a signal slightly above the background noise (P438: 46 detections/object, WT: 23 detections/object), PSynMyco one order of magnitude higher (104 detections/object) and PSpi giving the highest signal (858 detections/object).

**Figure 5:**
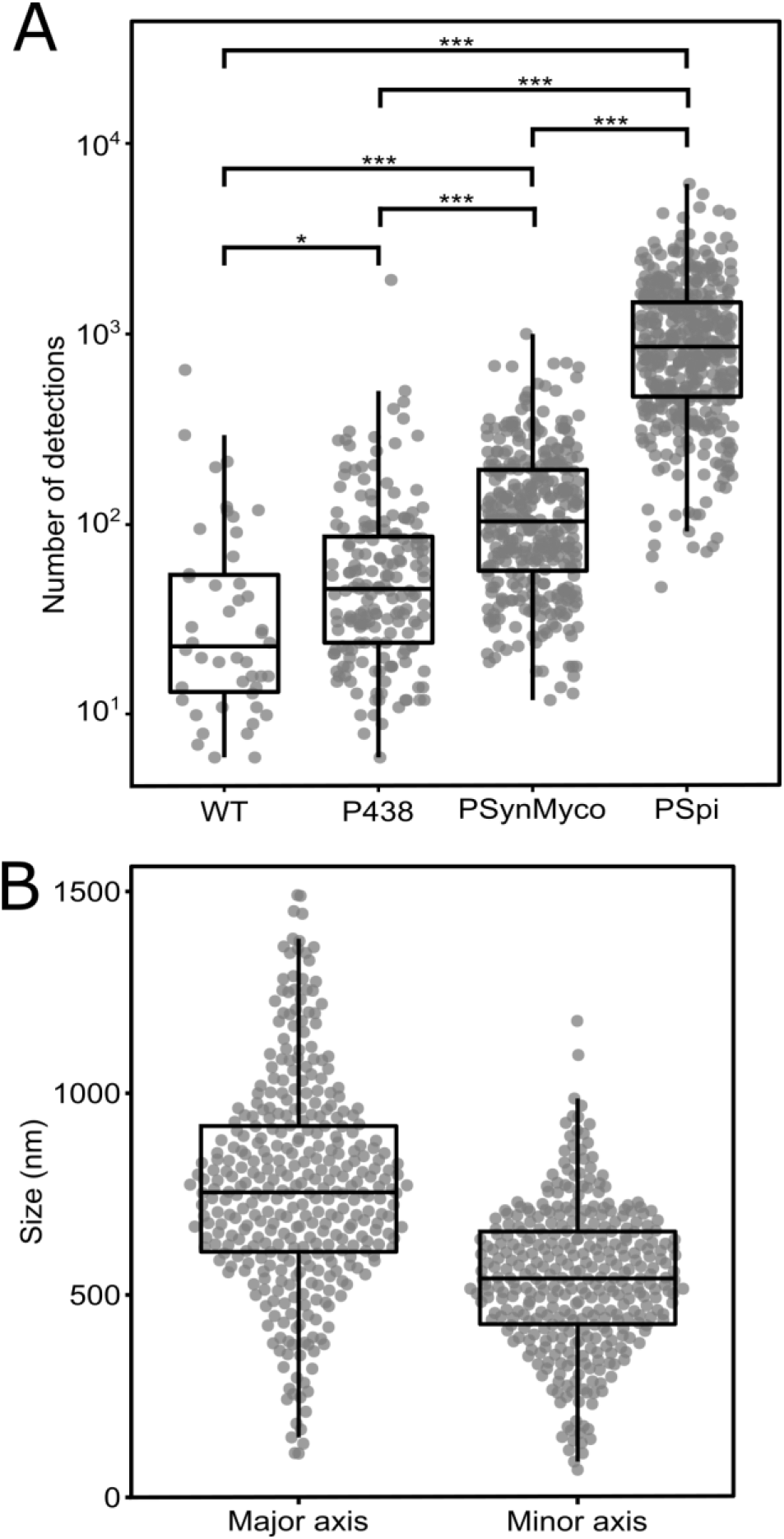
Evaluation of promoter strength and cell size through PALM imaging in *Mmc*. **A.** Comparison of promoter strength by PALM imaging. *Mmc* cells, either wild-type (WT) or transformed with the plasmids pMT85-P438-mEos3.2 (P438), pMT85-PSynMyco-mEos3.2 (PSynMyco) or pMT85-PSpi-mEos3.2 (PSpi) were imaged by PALM. For each strain, the data collected from a single representative field of view (512×512 pixels; pixel size: 0.16 μm) were analyzed. The dot-plot presents the number of detections measured in each object segmented by Tesseler (equivalent to a cell), on top of which a boxplot showing the median, interquartile range, minimum and maximum is overlaid. Statistics tests (Mann Whitney) were performed to compare the four strains (*: p-value <0.05; ***: p-value < 1.10^−10^). **B.** Deriving cell size data from PALM images. *Mmc* cells transformed with the plasmid pMT85-PSpi-mEos3.2 were imaged by PALM. The data collected from a single representative field of view (512×512 pixels; pixel size: 0.16 μm) were analyzed. The dot-plot presents the dimensions (in nm) of the major axis and the minor axis of each object segmented by Tesseler (Figure S7), on top of which a boxplot showing the median, interquartile range, minimum and maximum is overlaid.

Interestingly, the overexpression of mEos3.2 alone can also be used to estimate the absolute physical size of individual cells. Indeed, for each of the hundreds to thousands of cells found in the field of view, the cytoplasm can be reliably discriminated from the background based on the localization of the fluorescence signal. The surface occupied by the cytoplasm can be then accurately measured (length, width, area) (Figure S7). As an example, we analyzed a population of 358 cells of *Mmc* pMT85-PSpi-mEos3.2 (Figure 5B), showing a normal distribution of sizes ranging from 107 to 1490 nm (mean: 768 nm, median: 754 nm) on the major axis and 87 to 1179 nm (mean: 540 nm, median: 540 nm) on the minor axis.

## Discussion

Super-resolution microscopy techniques have revolutionized biology by enabling fluorescence imaging well beyond the diffraction limit. Establishing these methods in mycoplasmas represents a significant milestone for the investigation of these minute bacteria with atypical properties and broad biological relevance.

Here we report the first application of PALM and dSTORM, two mature SMLM techniques, in mycoplasmas. We describe a common, simple and reliable method to generate high-quality, SMLM-ready, fixed samples and showcase the interest of super-resolution imaging in our model species *Mycoplasma mycoides* subsp. *capri.* This bacterium, in addition to being the etiologic agent of Contagious Caprine PleuroPneumonia, has emerged as a major model in both synthetic and fundamental biology. Indeed, it is the original organism from which the synthetic cells JCVI Syn1.0 and Syn3.0 have been derived through genome engineering and reduction (12, 13). As a result, the SMLM strategies developed here in *Mmc* are likely to be applicable in its man-made descendants. We imaged two proteins of interest in *Mmc,* one cytoplasmic and one at the cell surface, with a pointing accuracy below 50 nm (more than 6 times the diffraction limit). The images reconstructed from our PALM and dSTORM datasets provide qualitative and quantitative data that were previously only accessible through immuno-gold electron microscopy. Our imaging process proved fast, as sample preparation takes ~3 hours for PALM and ~6 hours for dSTORM, followed by ~10-15 minutes of data acquisition for each field of view. Given that each field of view contains several hundred to several thousand cells, it is possible to collect data at a medium-throughput and to get an accurate description of a biological process across a large population of cells. In addition, we note that our process is highly reproducible, with similar results obtained across multiple fields of view acquired on the same coverslip, or across independent coverslips, prepared from independent biological replicates and imaged at different dates.

In this study, we focus our analysis on the localization of virulence factors associated with the cell membrane. However, SMLM can be applied to study any protein of interest, including those involved in fundamental biological processes such as transcription, translation or cell division, and has the potential to become a key tool for the mycoplasmologist community. In order to foster a quick adoption of SMLM, we have validated our sample preparation process in a total of six different *Mycoplasma* species, relevant to both the medical and veterinary fields and covering the three main *Mollicutes* phylogenetic subgroups.

We also provide the plasmid pMT85-PSynMyco-mEos3.2 which allows the expression of the photo-convertible fluorescent protein mEos3.2 under the control of the SynMyco regulatory element, enabling PALM imaging in all the tested *Mycoplasma* species. We then produced two variants of the plasmid in which the mEos3.2 expression is driven by the alternative promoters PSpi (51) or P438 (55), which are well known and used throughout the *Mollicutes* field. These three plasmids are available through the repository Addgene (references 173894, 173895, 173896) for rapid distribution, and evaluation in other species. It should be noted that the mEos3.2 is generally regarded as one of the best fluorophores for PALM imaging, as it exhibits multiple traits that are desirable for SMLM: it is monomeric, has a fast maturation and has a relatively high photostability. Its green and red excitation and emission wavelengths also make it compatible for multicolor imaging with widely used DNA dyes such as Hoechst 33342 (361/497 nm), membrane dyes such as Nile Red (552/636 nm) and far-red organic dyes such as Alexa 647 (650/670 nm). Alternative fluorescent proteins could be expressed from our pMT-85 backbone, and multiple options are available in the literature, including the well-characterized mMaple3 (79), PAmCherry (80) or Dendra2 (81). These alternatives should nonetheless be properly benchmarked, as it was shown the performances of different fluorescent proteins can vary greatly in a given microbial species (82).

Meanwhile, in order to perform dSTORM we relied on the expression of an epitope-tagged variant of our protein of interest which was subsequently immuno-labeled. This choice was guided by the possibility to use commercial, well-characterized, high affinity monoclonal anti-HA antibodies enabling us to have a good labeling of the protein of interest and a low background noise. However, genetic edition strategies allowing the precise modification of a genomic locus for epitope tagging are only available in a limited number of *Mycoplasma* species. This issue can however be bypassed by using custom, protein-specific antibodies. It is noteworthy that our protocol used a primary antibody and fluorescent-labeled secondary antibody. This method, while practical and widely used, introduces a significant displacement between the targeted protein and the reporting fluorophore due to the physical size of the antibodies (83). This linkage error is approximately of 15-20 nm and can be mitigated through the utilization of either a fluorophore-conjugated primary antibody or alternative small probes such as nanobodies or aptamers which offer a ~5 nm linkage error (84–86).

Finally, it should be noted that super-resolution microscopy is still a developing field, which sees continuous improvements in resolution through new techniques. For instance, strategies based around Expansion Microscopy (a process in which the sample is physically expanded in an isotropic fashion), could help reach the nanometer scale by decrowding biomolecules in dense samples (87). Meanwhile, the recently developed MINFLUX promises imaging at 1-3 nm lateral resolution, in multicolor and in 3D (88). These improvements, coupled with the commercialization of “turnkey” microscopy systems will further drive the adoption of SMLM and in the long term will benefit our understanding of mycoplasma’s biology.

## Author contribution

Funding acquisition: E.H. and Y.A.; Project administration: Y.A.; Supervision: E.H. and Y.A; Conceptualization: F.R., E.H. and Y.A; Resources: E.V. and E.H.; Investigation: F.R., A.V., P.B., E.H., and Y.A.; Formal analysis: F.R., E.H.; Visualization: F.R., E.H. and Y.A.; Writing: F.R., E.H. and Y.A.

## Acknowledgments

The authors would like to thank JB. Sibarita and Dr. F. Levet for providing access to the PALMTracer plugin and SR-Tesseler software, Dr. M. Mondin for her assistance during our first SMLM tests at the Bordeaux Imaging Center, Dr. T. Ipoutcha for his assistance with *M. gallisepticum* transformation, and Dr. N. Bourg and P. Barthelemy at Abbelight for fruitful discussion. We acknowledge C. Rouveyrol for her initial work on mycoplasma sample preparation. Y.A is personally indebted to C. Hunsa-Kredeinde for her kind review of an early draft of this manuscript. This study was funded by the French National Agency for Research (ANR) grant ANR-17-CE35-0002-01 DACSyMy.

---

**Figure S1:**
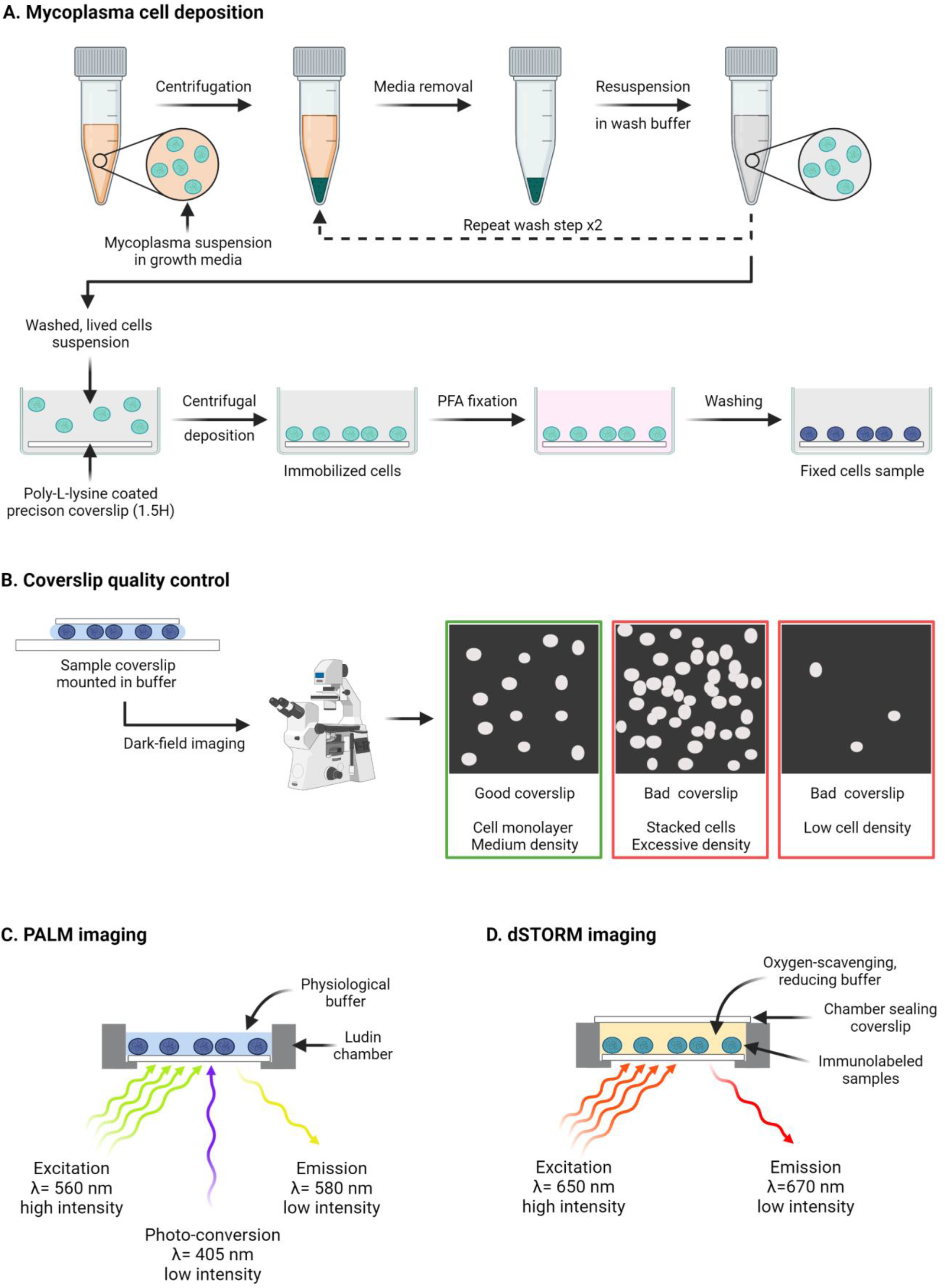
Schematic overview of the SMLM samples preparation process. **A.** Mycoplasma cells deposition and fixation on coverslips. Cells grown in culture media are harvested and washed, then deposited on poly-L-lysine coated coverslips using centrifugation to force the cells to sediment. After PFA fixation, the coverslips can be further processed by immune-labeling or used directly. **B.** Quality control of the sample coverslips. In order to check the proper deposition of cells on the coverslip, and to gauge the cell density, coverslips are mounted on glass slides (cells facing the glass) using a mounting buffer. Dark-field imaging is then performed to assess the samples. **C.** Schematic overview of the PALM imaging process. The coverslip is mounted in a Ludin chamber, filled with PBS buffer. Imaging is performed by providing continuous excitation illumination at medium intensity while simultaneously illuminating the sample with low-intensity ultra-violet light to photo-convert the fluorescent protein. **D.** Schematic overview of the dSTORM imaging process. The coverslip is mounted in dSTORM imaging buffer, in a sealed Ludin chamber. Imaging is performed by providing continuous excitation illumination at high intensity in order to send the fluorophores to their dark state.

**Figure S2.**
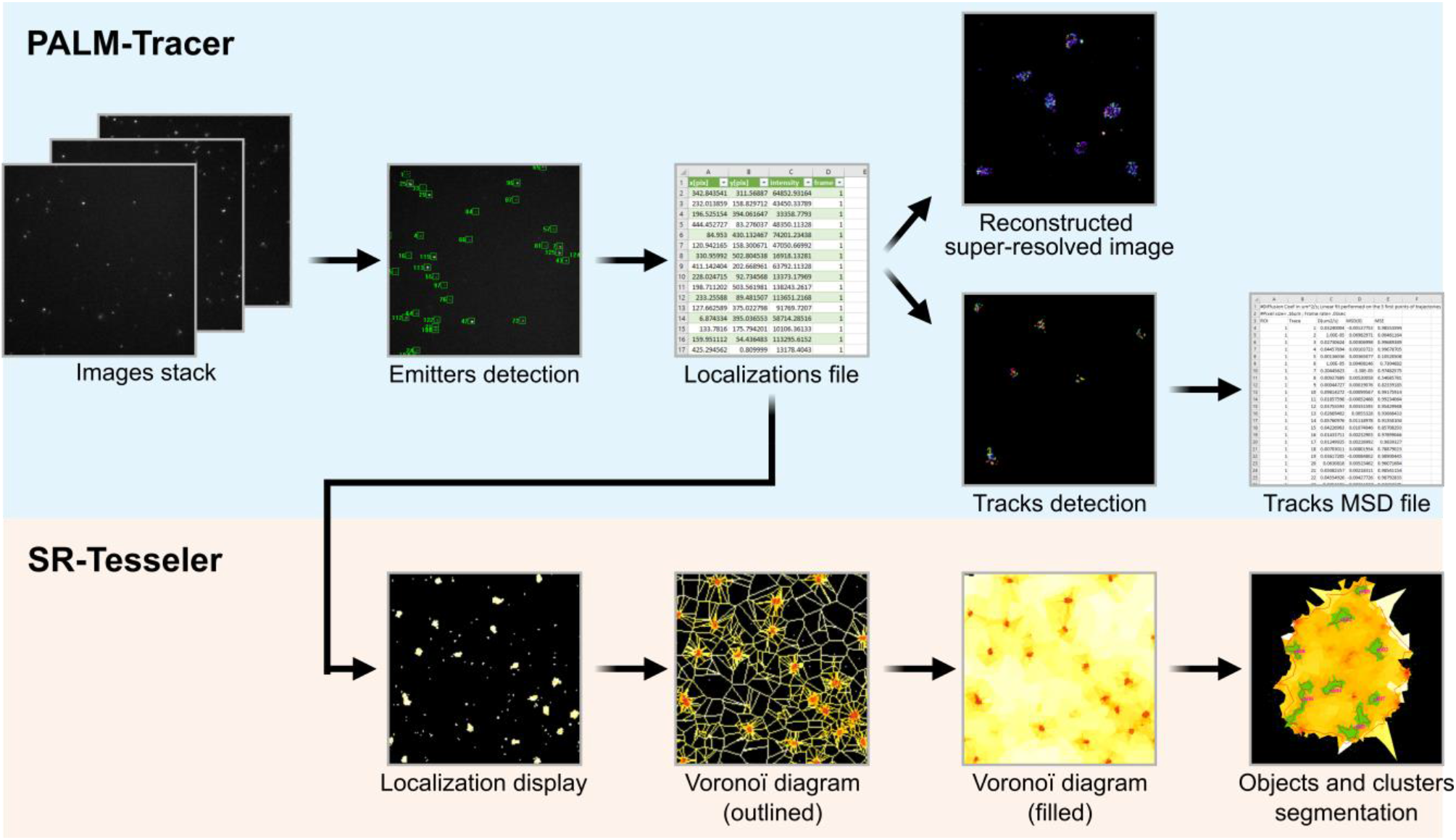
Schematic overview of the SMLM imaging data processing. Images stacks collected during data acquisition are processed using the Metamorph plug-in PALM-Tracer. Emitters are detected on each frame, and their localizations are computed by Gaussian fitting, yielding a localization file indicating for each emitter an x and a y localization value, an intensity value, and a frame number. This localization file is then used to reconstruct a super-resolved image, and to create emitters tracks from which the MSD values are derived. SR-Tesseler is used in parallel to analyze the localization file. First, a Voronoï diagram is computed, based on which objects (red outline) and clusters (green fill) are automatically segmented based on set density, detection and size threshold parameters.

**Figure S3:**
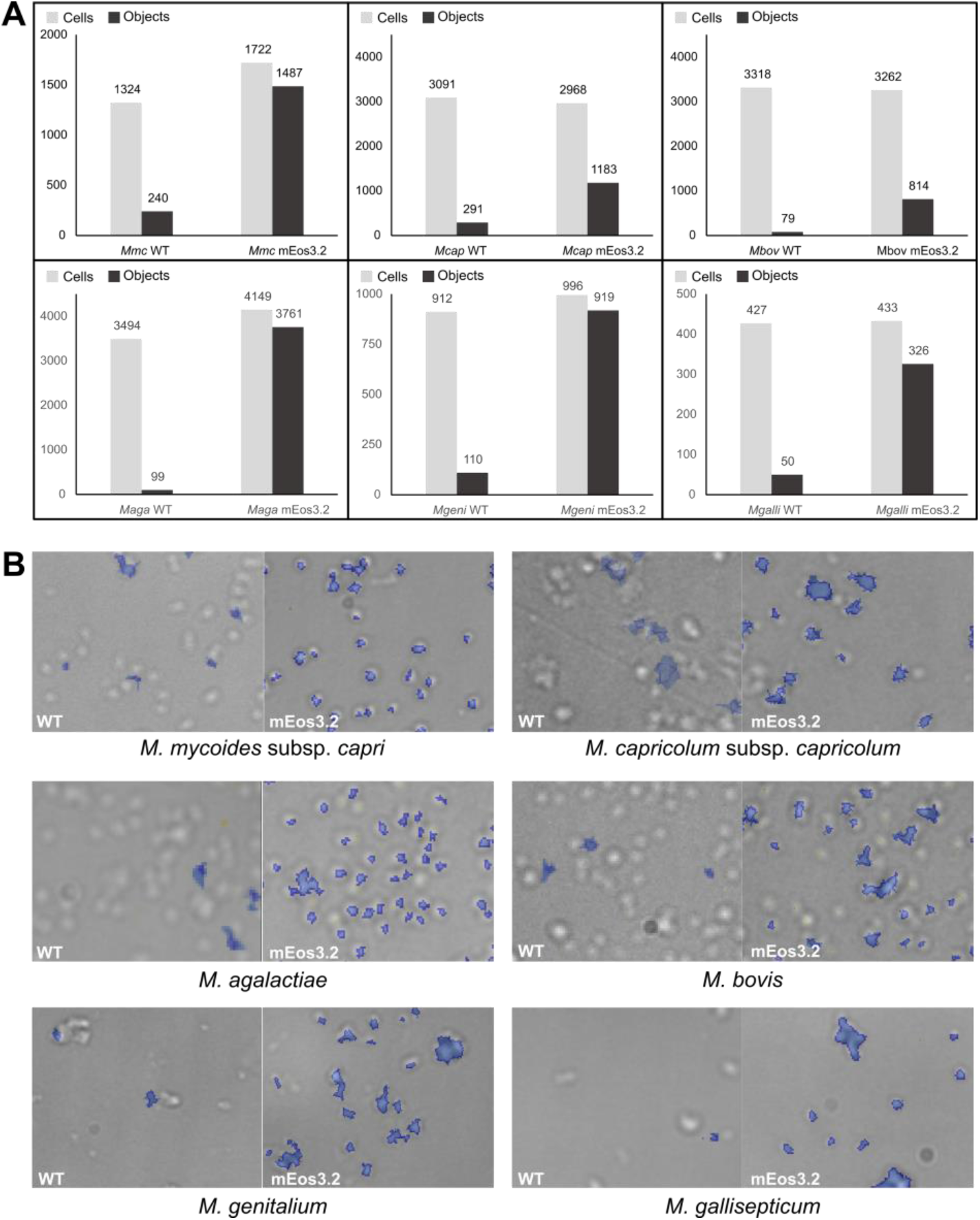
Assessing the functionality of the photo-convertible fluorescent protein mEos3.2 in multiple *Mycoplasma* species. Six mycoplasma species were transformed with the plasmid pMT85-PSynMyco-mEos3.2 and imaged by PALM (see Figure 1). The data presented here correspond to a single representative field of view (512×512 pixels; pixel size: 160 nm). **A.** Comparison of the number of cells and Tesseler segmented objects in the field of view. The bar graph indicates the number of objects segmented by Tesseler (equivalent to a cell) compared to the number of actual cells found in the imaged field. **B.** Localization of the Tesseler objects compared to the mycoplasma cells. Sample images show the overlay of a contrast-phase image and of the objects segmented by Tesseler (blue) in the same field.

**Figure S4:**
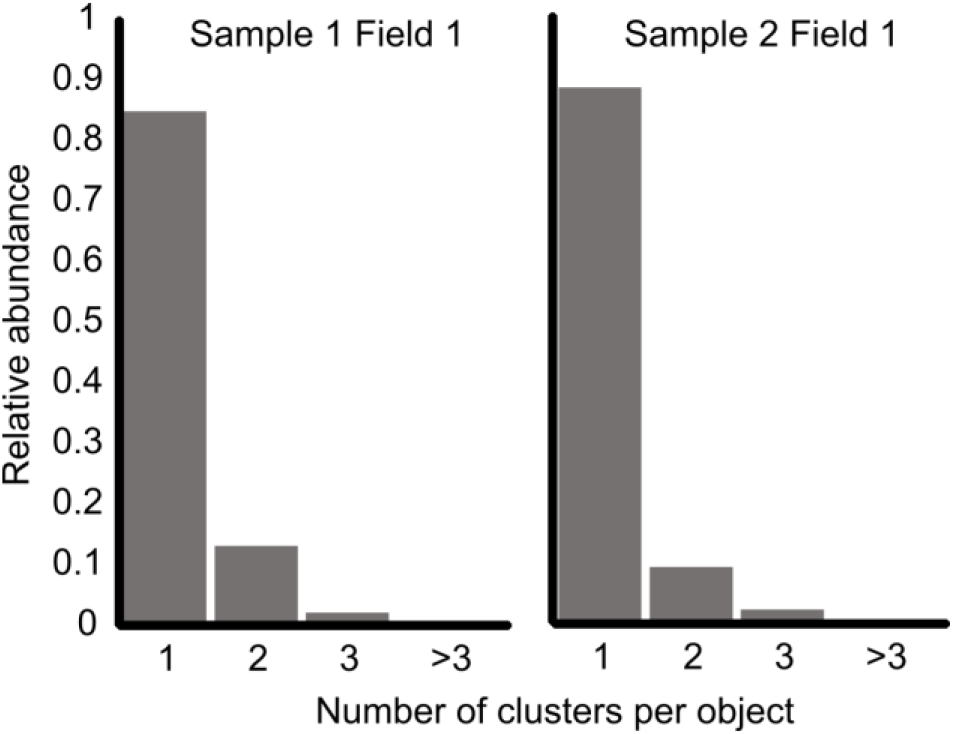
Tesseler clustering of mEos3.2 localizations in the cytoplasm of *Mycoplasma mycoides* subsp. *capri*. *Mmc* pMT85-PSynMyco-mEos3.2 cells were imaged using PALM, and Tesseler clustering of the fluorescence signal was performed. For each dataset, the number of clusters per Tesseler-segmented object was computed. The bar graphs display the distribution of the number of clusters per object. The data presented here correspond to two fields of view (512×512 pixels; pixel size: 160 nm), taken from two independent coverslips.

**Figure S5:**
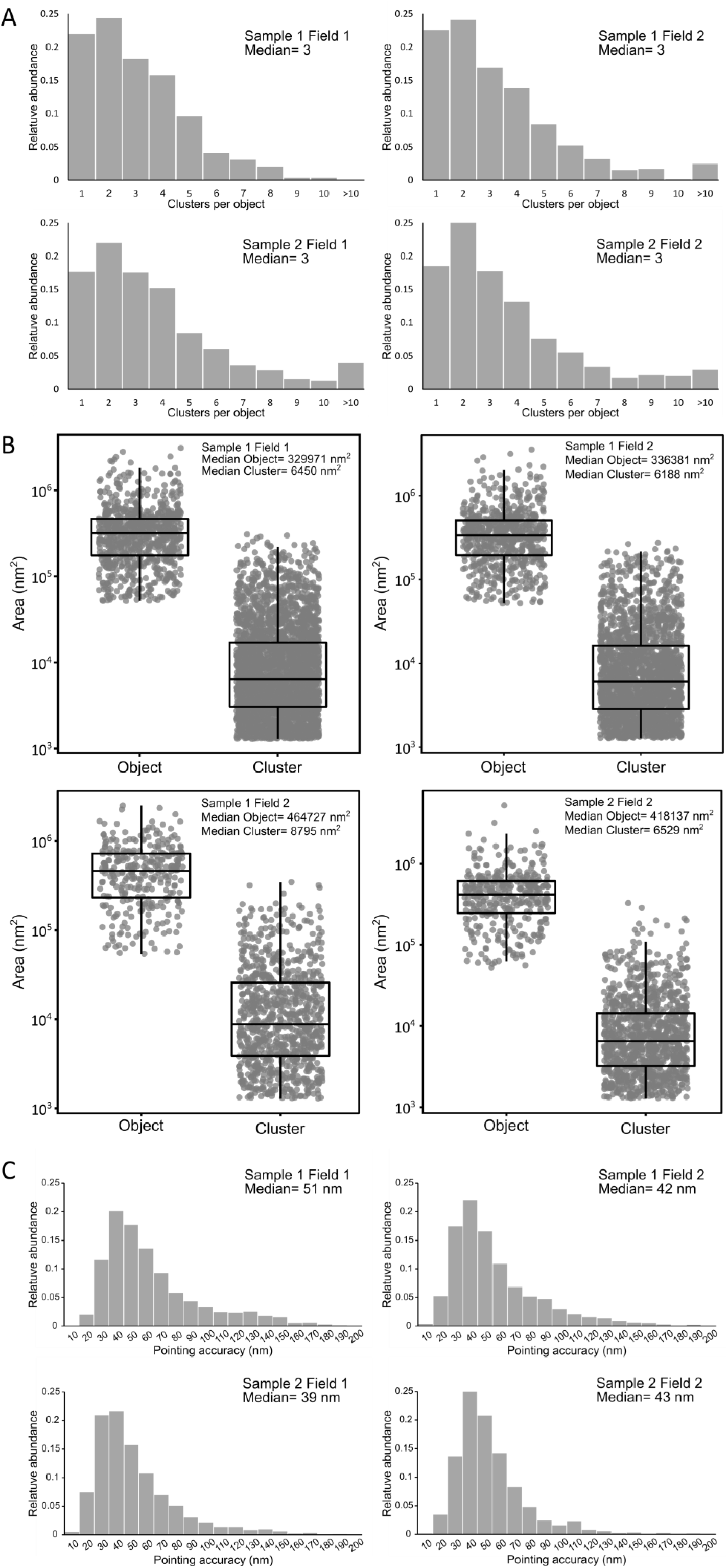
Replicates of the PALM imaging of an F-type ATPase in *Mycoplasma mycoides* subsp. *capri*. *Mmc* 0575-mEos3.2 cells, expressing a mEos-fused variant of the β-subunit of the ATPase F1-like domain, were imaged by PALM (see Figure 2). The data presented here correspond to four fields of view (512×512 pixels; pixel size: 160 nm) taken from two independent coverslips. **A.** Tesseler clustering of the fluorescence signal. The number of clusters per Tesseler-segmented objects was computed. The bar graphs display the distribution of the number of clusters per object. **B.** Objects and clusters sizes. The dot plot presents the area (in nm^2^) of each object and cluster segmented by Tesseler, to which a boxplot showing the median, interquartile range, minimum and maximum is overlaid. The median value of each data set is indicated. **B.** Evaluation of the PALM imaging pointing accuracy. The bar graphs display the distribution of the pointing accuracy derived from each track. The median value of each data set is indicated.

**Figure S6:**
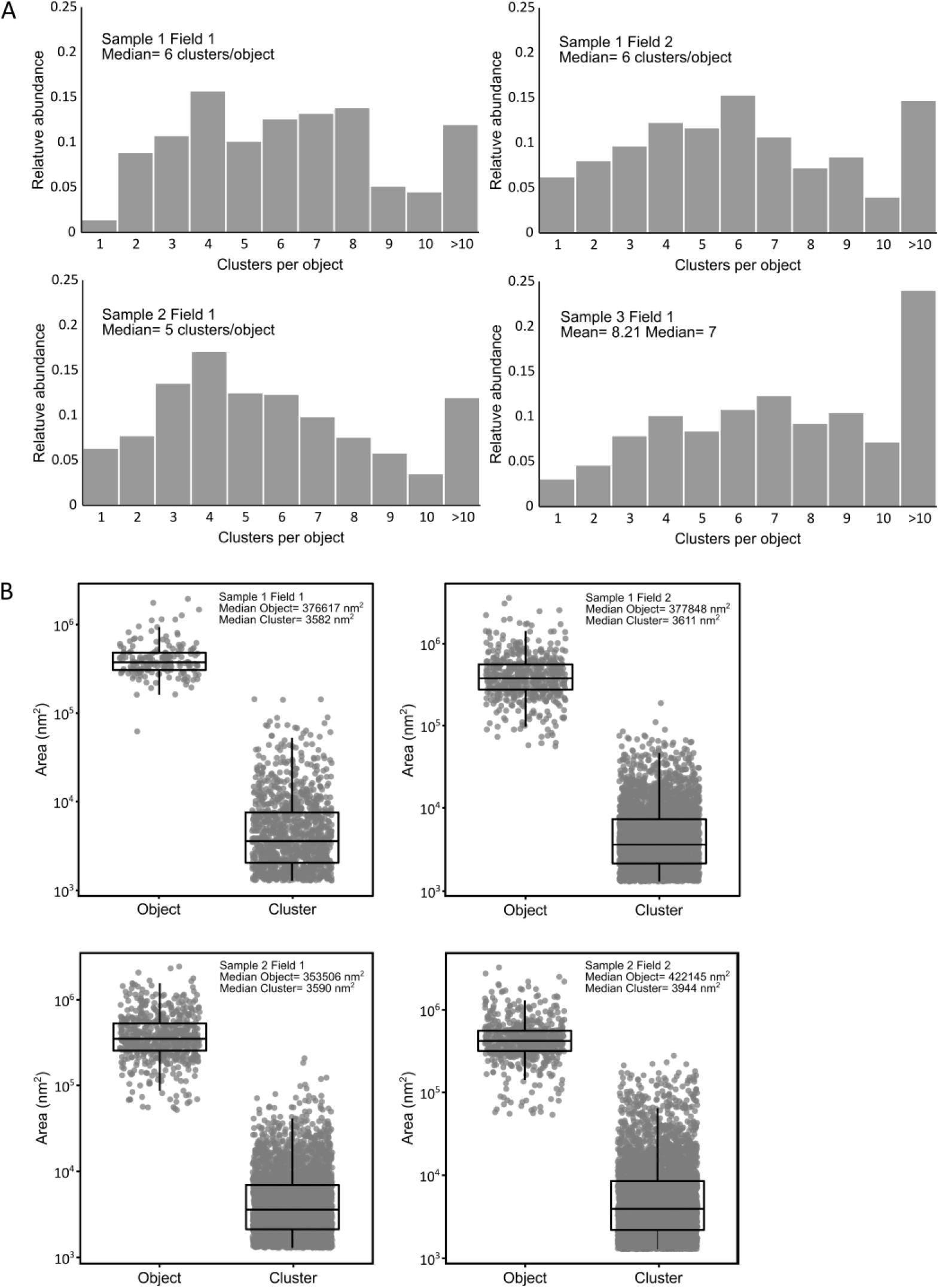
Replicates of the dSTORM imaging of an antibody-specific protease in *Mycoplasma mycoides* subsp. *capri*. *Mmc* 0582-HA cells expressing an HA-tag fused variant of the serine protease MIP0582 were immune-labeled and imaged by dSTORM (see Figure 3). The data presented here correspond to four fields of view (512×512 pixels; pixel size: 160 nm) taken from two independent coverslips. **A.** Tesseler clustering of the fluorescence signal. For each field of view, the number of clusters per Tesseler-segmented objects was computed. The bar graphs display the distribution of the number of clusters per object. **B.** Objects and clusters sizes. The dot plot presents the area (in nm^2^) of each object and cluster segmented by Tesseler, to which a boxplot showing the median, interquartile range, minimum and maximum is overlaid. The median value of each data set is indicated.

**Figure S7:**
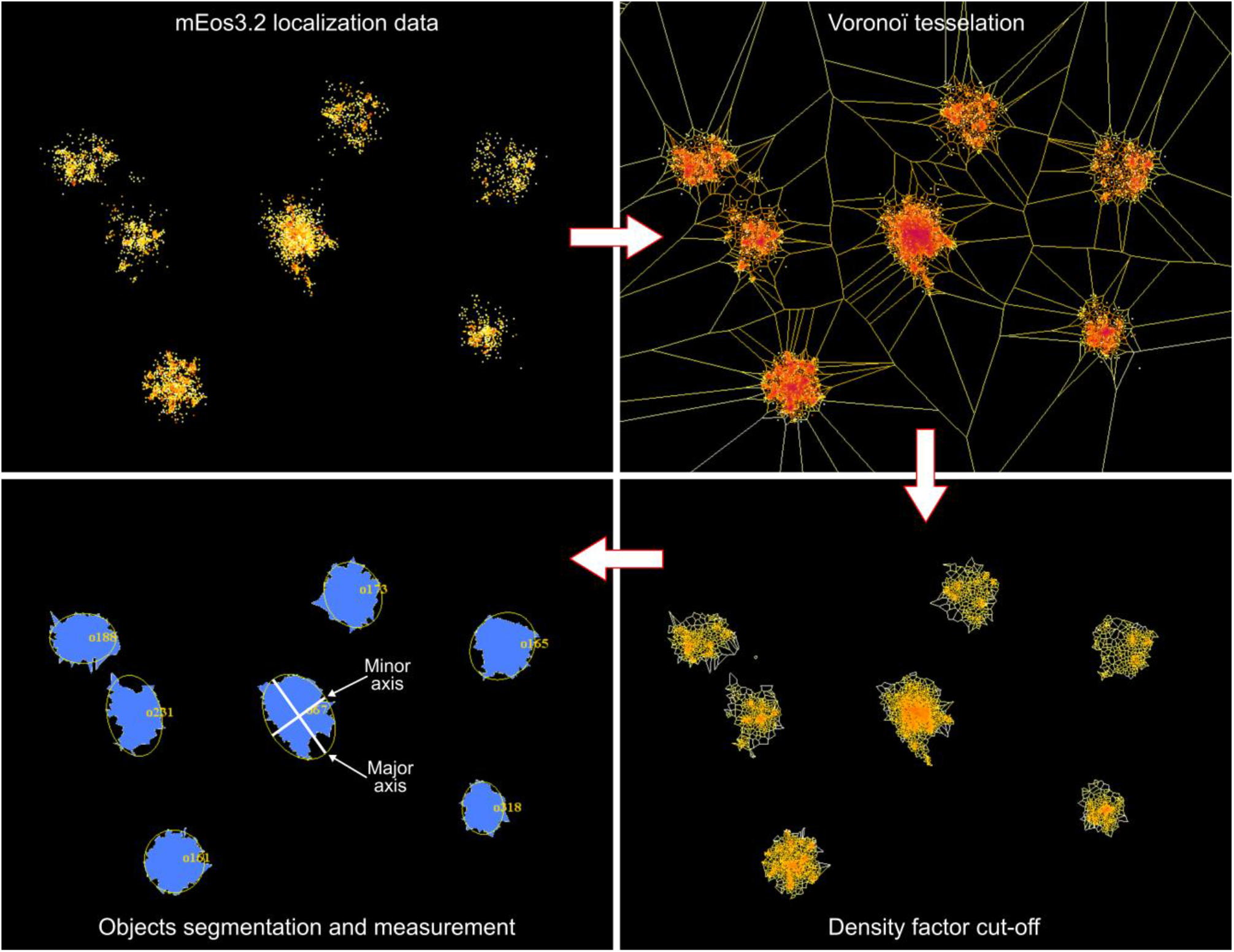
Schematic overview of the cell size estimation from PALM imaging data. Top-left: localization data collected from *Mmc* cells expressing the mEos3.2 as a soluble monomer in the cytoplasm are displayed in Tesseler. Top-right: Voronoï tessellation of the localizations (no density cut-off). Bottom-right: Voronoï tessellation of the localizations is performed again after setting a density factor cut-off (here: 1) in order to filter the signal coming from outside the cells. Bottom-left: Tesseler objects are segmented and their dimensions (major axis and minor axis) are computed.

## Notes

### Competing Interest Statement

The authors have declared no competing interest.

## References

1. P. Sirand-Pugnet, C. Citti, A. Barré, A. Blanchard, Evolution of mollicutes: down a bumpy road with twists and turns. Res. Microbiol. 158, 754–766 (2007).

2. S. Razin, “The Genus Mycoplasma and Related Genera (Class Mollicutes)” in The Prokaryotes, (Springer US, 2006), pp. 836–904.

3. M. May, M. F. Balish, A. Blanchard, “The Order Mycoplasmatales” in The Prokaryotes, (Springer Berlin Heidelberg, 2014), pp. 515–550.

4. C. M. Fraser, et al., The minimal gene complement of Mycoplasma genitalium. Science 270, 397–403 (1995).

5. P. Schwille, et al., MaxSynBio: Avenues Towards Creating Cells from the Bottom Up. Angew. Chem. Int. Ed. Engl. 57, 13382–13392 (2018).

6. M. Juhas, L. Eberl, J. I. Glass, Essence of life: essential genes of minimal genomes. Trends Cell Biol. 21, 562–8 (2011).

7. M. Lluch-Senar, E. Querol, J. Piñol, Cell division in a minimal bacterium in the absence of ftsZ. Mol. Microbiol. 78, 278–89 (2010).

8. C. A. Hutchison, et al., Global transposon mutagenesis and a minimal Mycoplasma genome. Science 286, 2165–9 (1999).

9. G. Y. Fisunov, et al., Core proteome of the minimal cell: comparative proteomics of three mollicute species. PLoS One 6, e21964 (2011).

10. M. Lluch-Senar, et al., Defining a minimal cell: essentiality of small ORFs and ncRNAs in a genome-reduced bacterium. Mol. Syst. Biol. 11, 780 (2015).

11. C. Henry, R. Overbeek, R. L. Stevens, Building the blueprint of life. Biotechnol. J. 5, 695–704 (2010).

12. D. G. G. Gibson, et al., Creation of a bacterial cell controlled by a chemically synthesized genome. Science 329, 52–6 (2010).

13. C. A. Hutchison, et al., Design and synthesis of a minimal bacterial genome. Science 351, aad6253 (2016).

14. J. R. Karr, et al., A whole-cell computational model predicts phenotype from genotype. Cell 150, 389–401 (2012).

15. M. Maritan, et al., Building Structural Models of a Whole Mycoplasma Cell. J. Mol. Biol. 434, 167351 (2022).

16. Z. R. Thornburg, et al., Fundamental behaviors emerge from simulations of a living minimal cell. Cell 185, 345–360.e28 (2022).

17. D. Taylor-Robinson, J. S. Jensen, Mycoplasma genitalium: from Chrysalis to multicolored butterfly. Clin. Microbiol. Rev. 24, 498–514 (2011).

18. F. M. Sánchez-Vargas, O. G. Gómez-Duarte, Mycoplasma pneumoniae-an emerging extra-pulmonary pathogen. Clin. Microbiol. Infect. 14, 105–17 (2008).

19. E. Biondi, et al., Treatment of mycoplasma pneumonia: a systematic review. Pediatrics 133, 1081–90 (2014).

20. H. Moi, K. Blee, P. J. Horner, Management of non-gonococcal urethritis. BMC Infect. Dis. 15, 294 (2015).

21. R. A. J. A. J. Nicholas, R. D. D. Ayling, Mycoplasma bovis: disease, diagnosis, and control. Res. Vet. Sci. 74, 105–112 (2003).

22. G. Bölske, et al., Diagnosis of contagious caprine pleuropneumonia by detection and identification of Mycoplasma capricolum subsp. capripneumoniae by PCR and restriction enzyme analysis. J. Clin. Microbiol. 34, 785–91 (1996).

23. D. Maes, et al., Update on Mycoplasma hyopneumoniae infections in pigs: Knowledge gaps for improved disease control. Transbound. Emerg. Dis. 65 Suppl 1, 110–124 (2018).

24. J. D. Evans, et al., Mycoplasma gallisepticum: Current and developing means to control the avian pathogen. J. Appl. Poult. Res. 14, 757–763 (2005).

25. I. Tsarmpopoulos, et al., In-Yeast Engineering of a Bacterial Genome Using CRISPR/Cas9. ACS Synth Biol 5, 104–109 (2016).

26. G. Christiansen, S. Birkelund, Transmission electron microscopy and immunogold staining of mollicute surface antigens. Methods Mol. Biol. 104, 309–18 (1998).

27. M. K. Stevens, D. C. Krause, Mycoplasma pneumoniae cytadherence phase-variable protein HMW3 is a component of the attachment organelle. J. Bacteriol. 174, 4265–4274 (1992).

28. M. K. Stevens, D. C. Krause, Localization of the Mycoplasma pneumoniae cytadherence-accessory proteins HMW1 and HMW4 in the cytoskeletonlike Triton shell. J. Bacteriol. 173, 1041–50 (1991).

29. O. Neyrolles, et al., Phase variations of the Mycoplasma penetrans main surface lipoprotein increase antigenic diversity. Infect. Immun. 67, 1569–78 (1999).

30. M. Schumacher, et al., Evidence for the Cytoplasmic Localization of the L-α-Glycerophosphate Oxidase in Members of the “Mycoplasma mycoides Cluster.” Front. Microbiol. 10, 1–13 (2019).

31. S. Seto, G. Layh-Schmitt, T. Kenri, M. Miyata, Visualization of the Attachment Organelle and Cytadherence Proteins of Mycoplasma pneumoniae by Immunofluorescence Microscopy. J. Bacteriol. 183, 1621–1630 (2001).

32. S. Seto, M. Miyata, Attachment organelle formation represented by localization of cytadherence proteins and formation of the electron-dense core in wild-type and mutant strains of Mycoplasma pneumoniae. J. Bacteriol. 185, 1082–1091 (2003).

33. D. Nakane, M. Miyata, Cytoskeletal “jellyfish” structure of Mycoplasma mobile. Proc. Natl. Acad. Sci. U. S. A. 104, 19518–19523 (2007).

34. H. N. Wu, M. Miyata, Whole surface image of Mycoplasma mobile, suggested by protein identification and immunofluorescence microscopy. J. Bacteriol. 194, 5848–55 (2012).

35. T. Kenri, et al., Use of Fluorescent-Protein Tagging To Determine the Subcellular Localization of Mycoplasma pneumoniae Proteins Encoded by the Cytadherence Regulatory Locus. 186, 6944–6955 (2004).

36. C. Martínez-Torró, et al., Functional Characterization of the Cell Division Gene Cluster of the Wall-less Bacterium Mycoplasma genitalium. Front. Microbiol. 12, 695572 (2021).

37. B. M. Hasselbring, J. L. Jordan, R. W. Krause, D. C. Krause, Terminal organelle development in the cell wall-less bacterium Mycoplasma pneumoniae. Proc. Natl. Acad. Sci. U. S. A. 103, 16478–16483 (2006).

38. I. Tulum, K. Kimura, M. Miyata, Identification and sequence analyses of the gliding machinery proteins from Mycoplasma mobile. Sci. Rep. 10, 3792 (2020).

39. D. Nakane, T. Kenri, L. Matsuo, M. Miyata, Systematic Structural Analyses of Attachment Organelle in Mycoplasma pneumoniae. PLOS Pathog. 11, e1005299 (2015).

40. H. Z. A. Ishag, et al., GFP as a marker for transient gene transfer and expression in Mycoplasma hyorhinis. Springerplus 5, 769 (2016).

41. A. Montero-Blay, S. Miravet-Verde, M. Lluch-Senar, C. Piñero-Lambea, L. Serrano, SynMyco transposon: engineering transposon vectors for efficient transformation of minimal genomes. DNA Res. 26, 327–339 (2019).

42. T. Bonnefois, et al., Development of fluorescence expression tools to study host-mycoplasma interactions and validation in two distant mycoplasma clades. J. Biotechnol. 236, 35–44 (2016).

43. L. Schermelleh, R. Heintzmann, H. Leonhardt, A guide to super-resolution fluorescence microscopy. J. Cell Biol. 190, 165–175 (2010).

44. B. Turkowyd, D. Virant, U. Endesfelder, From single molecules to life: microscopy at the nanoscale. Anal. Bioanal. Chem. 408, 6885–6911 (2016).

45. B. O. Leung, K. C. Chou, Review of Super-Resolution Fluorescence Microscopy for Biology. Appl. Spectrosc. 65, 967–980 (2011).

46. M. Lelek, et al., Single-molecule localization microscopy. Nat. Rev. Methods Prim. 1 (2021).

47. M. Sauer, M. Heilemann, Single-Molecule Localization Microscopy in Eukaryotes. Chem. Rev. 117, 7478–7509 (2017).

48. H. H. Tuson, J. S. Biteen, Unveiling the inner workings of live bacteria using super-resolution microscopy. Anal. Chem. 87, 42–63 (2015).

49. S. Holden, Probing the mechanistic principles of bacterial cell division with super-resolution microscopy. Curr. Opin. Microbiol. 43, 84–91 (2018).

50. A. Gahlmann, W. E. Moerner, Exploring bacterial cell biology with single-molecule tracking and super-resolution imaging. Nat. Rev. Microbiol. 12, 9–22 (2014).

51. F. Labroussaa, et al., Impact of donor–recipient phylogenetic distance on bacterial genome transplantation. Nucleic Acids Res. 44, 8501–8511 (2016).

52. E. A. Freundt, “CULTURE MEDIA FOR CLASSIC MYCOPLASMAS” in Methods in Mycoplasmology, (Elsevier, 1983), pp. 127–135.

53. J. G. Tully, D. L. Rose, R. F. Whitcomb, R. P. Wenzel, Enhanced isolation of Mycoplasma pneumoniae from throat washings with a newly-modified culture medium. J. Infect. Dis. 139, 478–82 (1979).

54. A. Montero-Blay, S. Miravet-Verde, M. Lluch-Senar, C. Piñero-Lambea, L. Serrano, SynMyco transposon: engineering transposon vectors for efficient transformation of minimal genomes. DNA Res. (2019) https:/doi.org/10.1093/dnares/dsz012.

55. O. Q. Pich, R. Burgos, R. Planell, E. Querol, J. Piñol, Comparative analysis of antibiotic resistance gene markers in Mycoplasma genitalium: application to studies of the minimal gene complement. Microbiology 152, 519–527 (2006).

56. K. W. King, K. Dybvig, Transformation of Mycoplasma capricolum and examination of DNA restriction modification in M. capricolum and Mycoplasma mycoides subsp. mycoides. Plasmid 31, 308–311 (1994).

57. X. Zhu, et al., Mbov_0503 Encodes a Novel Cytoadhesin that Facilitates Mycoplasma bovis Interaction with Tight Junctions. Microorganisms 8(2020).

58. S. P. Reddy, W. G. Rasmussen, J. B. Baseman, Isolation and characterization of transposon Tn *4001*-generated, cytadherence-deficient transformants of *Mycoplasma pneumoniaeand Mycoplasma genitalium*. FEMS Immunol. Med. Microbiol. 15, 199–211 (1996).

59. P. Nottelet, et al., The mycoplasma surface proteins MIB and MIP promote the dissociation of the antibody-antigen interaction. Sci. Adv. 7, eabf2403 (2021).

60. D. Nair, et al., Super-resolution imaging reveals that AMPA receptors inside synapses are dynamically organized in nanodomains regulated by PSD95. J. Neurosci. 33, 13204–24 (2013).

61. A. Kechkar, D. Nair, M. Heilemann, D. Choquet, J.-B. Sibarita, Real-time analysis and visualization for single-molecule based super-resolution microscopy. PLoS One 8, e62918 (2013).

62. I. Izeddin, et al., Wavelet analysis for single molecule localization microscopy. Opt. Express 20, 2081–95 (2012).

63. X. Michalet, Mean square displacement analysis of single-particle trajectories with localization error: Brownian motion in an isotropic medium. Phys. Rev. E 82, 041914 (2010).

64. F. Levet, et al., SR-Tesseler: a method to segment and quantify localization-based super-resolution microscopy data. Nat. Methods 12, 1065–71 (2015).

65. M. Postma, J. Goedhart, PlotsOfData-A web app for visualizing data together with their summaries. PLoS Biol. 17, e3000202 (2019).

66. A. R. Halpern, M. D. Howard, J. C. Vaughan, Point by Point: An Introductory Guide to Sample Preparation for Single-Molecule, Super-Resolution Fluorescence Microscopy. Curr. Protoc. Chem. Biol. 7, 103–20 (2015).

67. A. Jimenez, K. Friedl, C. Leterrier, About samples, giving examples: Optimized Single Molecule Localization Microscopy. Methods 174, 100–114 (2020).

68. E. A. Rodriguez, et al., The Growing and Glowing Toolbox of Fluorescent and Photoactive Proteins. Trends Biochem. Sci. 42, 111–129 (2017).

69. M. Zhang, et al., Rational design of true monomeric and bright photoactivatable fluorescent proteins. Nat. Methods 9, 727–729 (2012).

70. C.-U. Zimmerman, R. Herrmann, Synthesis of a small, cysteine-rich, 29 amino acids long peptide in *Mycoplasma pneumoniae*. FEMS Microbiol. Lett. 253, 315–321 (2005).

71. E. Dordet Frisoni, et al., ICEA of Mycoplasma agalactiae: a new family of self-transmissible integrative elements that confers conjugative properties to the recipient strain. Mol. Microbiol. 89, 1226–1239 (2013).

72. M. Breton, et al., First report of a tetracycline-inducible gene expression system for mollicutes. Microbiology 156, 198–205 (2010).

73. C. Janis, et al., Unmarked insertional mutagenesis in the bovine pathogen Mycoplasma mycoides subsp. mycoides SC: characterization of a lppQ mutant. Microbiology 154, 2427–2436 (2008).

74. F. Rideau, et al., Random transposon insertion in the Mycoplasma hominis minimal genome. Sci. Rep. 9, 13554 (2019).

75. B. R. Lyon, J. W. May, R. A. Skurray, Tn4001: a gentamicin and kanamycin resistance transposon in Staphylococcus aureus. Mol. Gen. Genet. 193, 554–556 (1984).

76. L. Béven, et al., Specific evolution of F1-like ATPases in mycoplasmas. PLoS One 7, e38793 (2012).

77. R. J. Davey, M. A. Digman, E. Gratton, P. D. J. Moens, Quantitative image mean squared displacement (iMSD) analysis of the dynamics of profilin 1 at the membrane of live cells. Methods 140–141, 119–125 (2018).

78. Y. Arfi, et al., MIB-MIP is a mycoplasma system that captures and cleaves immunoglobulin G. Proc. Natl. Acad. Sci. U. S. A. 113, 1600546113- (2016).

79. S. Wang, J. R. Moffitt, G. T. Dempsey, X. S. Xie, X. Zhuang, Characterization and development of photoactivatable fluorescent proteins for single-molecule-based superresolution imaging. Proc. Natl. Acad. Sci. U. S. A. 111, 8452–7 (2014).

80. F. V Subach, et al., Photoactivatable mCherry for high-resolution two-color fluorescence microscopy. Nat. Methods 6, 153–9 (2009).

81. N. G. Gurskaya, et al., Engineering of a monomeric green-to-red photoactivatable fluorescent protein induced by blue light. Nat. Biotechnol. 24, 461–5 (2006).

82. B. Turkowyd, et al., Establishing Live-Cell Single-Molecule Localization Microscopy Imaging and Single-Particle Tracking in the Archaeon Haloferax volcanii. Front. Microbiol. 11 (2020).

83. S. M. Früh, et al., Site-Specifically-Labeled Antibodies for Super-Resolution Microscopy Reveal In Situ Linkage Errors. ACS Nano (2021) https:/doi.org/10.1021/acsnano.1c03677.

84. S. Sograte-Idrissi, et al., Nanobody Detection of Standard Fluorescent Proteins Enables Multi-Target DNA-PAINT with High Resolution and Minimal Displacement Errors. Cells 8 (2019).

85. J. Ries, C. Kaplan, E. Platonova, H. Eghlidi, H. Ewers, A simple, versatile method for GFP-based super-resolution microscopy via nanobodies. Nat. Methods 9, 582–584 (2012).

86. R. P. Moore, W. R. Legant, Improving probes for super-resolution. Nat. Methods 15, 659–660 (2018).

87. A. T. Wassie, Y. Zhao, E. S. Boyden, Expansion microscopy: principles and uses in biological research. Nat. Methods 16, 33–41 (2019).

88. K. C. Gwosch, et al., MINFLUX nanoscopy delivers 3D multicolor nanometer resolution in cells. Nat. Methods 17, 217–224 (2020).

